# High-performance brain-to-text communication via imagined handwriting

**DOI:** 10.1101/2020.07.01.183384

**Authors:** Francis R. Willett, Donald T. Avansino, Leigh R. Hochberg, Jaimie M. Henderson, Krishna V. Shenoy

## Abstract

Brain-computer interfaces (BCIs) can restore communication to people who have lost the ability to move or speak. To date, a major focus of BCI research has been on restoring gross motor skills, such as reaching and grasping^1–5^ or point-and-click typing with a 2D computer cursor^6,7^. However, rapid sequences of highly dexterous behaviors, such as handwriting or touch typing, might enable faster communication rates. Here, we demonstrate an intracortical BCI that can decode imagined handwriting movements from neural activity in motor cortex and translate it to text in real-time, using a novel recurrent neural network decoding approach. With this BCI, our study participant (whose hand was paralyzed) achieved typing speeds that exceed those of any other BCI yet reported: 90 characters per minute at >99% accuracy with a general-purpose autocorrect. These speeds are comparable to able-bodied smartphone typing speeds in our participant’s age group (115 characters per minute)^8^ and significantly close the gap between BCI-enabled typing and able-bodied typing rates. Finally, new theoretical considerations explain why temporally complex movements, such as handwriting, may be fundamentally easier to decode than point-to-point movements. Our results open a new approach for BCIs and demonstrate the feasibility of accurately decoding rapid, dexterous movements years after paralysis.

## Results

Prior BCI studies have shown that the motor intention for gross motor skills, such as reaching, grasping or moving a computer cursor, remains neurally encoded in motor cortex after paralysis^1–7^. However, it is still unknown whether the neural representation for a rapid and highly-dexterous motor skill, such as handwriting, also remains intact. We tested this by recording neural activity from two microelectrode arrays in the hand “knob” area of precentral gyrus^9,10^ while our study participant, T5, attempted to handwrite individual letters and symbols (Fig. 1A). T5 has a high-level spinal cord injury and was paralyzed from the neck down. We instructed T5 to “attempt” to write as if his hand was not paralyzed (while imagining that he was holding a pen on a piece of ruled paper).

**Figure 1.**
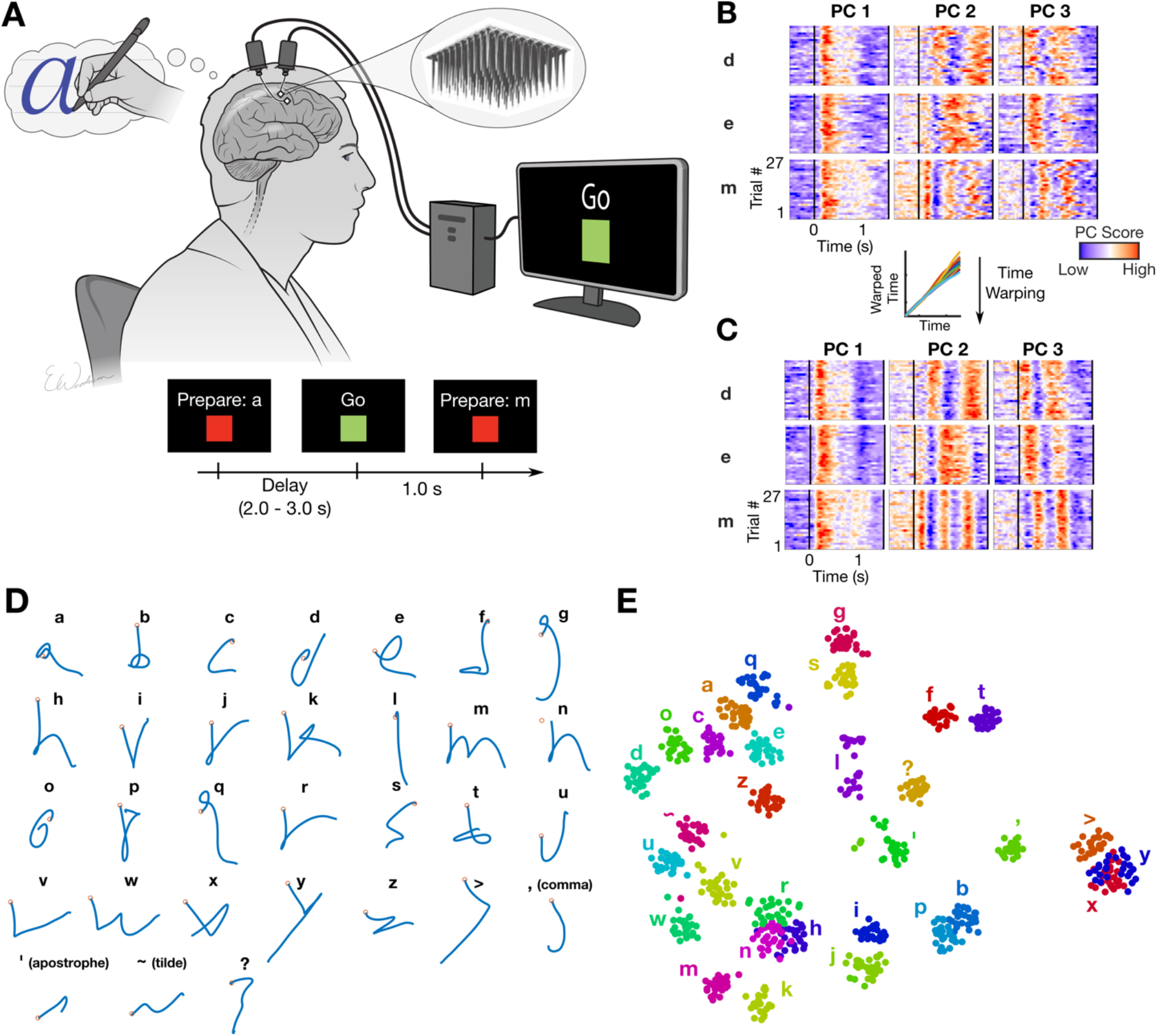
Robust neural encoding of attempted handwriting. **(A)** Participant T5 attempted to handwrite each character one at a time, following the instructions given on a computer screen (lower panels depict what is shown on the screen, following the timeline). **(B)** Neural activity in the top 3 principal components (PCs) is shown for three example letters (d, e and m) and 27 repetitions of each letter (“trials”). The color scale was normalized within each panel separately for visualization. **(C)** Time-warping the neural activity to remove trial-to-trial changes in writing speed reveals consistent patterns of activity unique to each letter. In the inset above C, example time-warping functions are shown for the letter “m”, and lie relatively close to the identity line (each trial’s warping function is plotted with a differently colored line). **(D)** Decoded pen trajectories are shown for all 31 tested characters: 26 lower-case letters, commas, apostrophes, question marks, tildes (~) and greater-than signs (>). Intended 2D pen tip velocity was linearly decoded from the neural activity using cross-validation (each character was held out). The decoded velocity was then averaged across trials and integrated to compute the pen trajectory (orange circles denote the start of the trajectory). **(E)** A 2-dimensional visualization of the neural activity made using t-SNE. Each circle is a single trial (27 trials for each of 31 characters).

We used principal components analysis to reduce the recorded neural activity (multiunit threshold crossing rates) to the top 3 dimensions containing the most variance (Fig. 1B). The neural activity appeared to be strong and repeatable, although the timing of its peaks and valleys varied across trials (potentially due to fluctuations in writing speed). We used a time-alignment technique to remove temporal variability^11^, revealing remarkably consistent underlying patterns of neural activity that are unique to each character (Fig. 1C). To see if the neural activity encoded the pen movements needed to draw each character’s shape, we attempted to reconstruct each character by linearly decoding pen tip velocity from the neural activity (Fig. 1D). Readily recognizable letter shapes confirm that pen tip velocity is robustly encoded. Finally, we used a nonlinear dimensionality reduction method (t-SNE) to produce a 2-dimensional visualization of each trial’s neural activity recorded after the “go” cue was given (Fig. 1E). The t-SNE visualization revealed tight clusters of neural activity for each character and a predominantly motoric encoding (where characters that are written similarly are closer together). Using a k-nearest neighbor classifier applied to the neural activity, we could classify the characters with 94.1% accuracy (chance level = 3.2%). Taken together, these results suggest that, even years after paralysis, the neural representation of handwriting in motor cortex is likely strong enough to be useful for a BCI.

Next, we tested whether we could decode complete handwritten sentences in real-time, thus enabling someone with paralysis to communicate by attempting to handwrite their intended message. To do so, we developed specialized methods to train a recurrent neural network (RNN) to convert the neural activity into probabilities describing the likelihood of each character being written at each moment in time (Fig. 2A, SFig. 1). These probabilities could either be thresholded in a simple way to emit discrete characters, which we did for real-time decoding (Fig. 2A “Raw Output”), or processed more extensively by a language model to simulate an autocorrect feature, which we applied retrospectively (Fig. 2A “Retrospective Output from a Language Model”). We used the limited set of 31 characters shown in Fig. 1D, consisting of the 26 lower case letters of the alphabet, commas, apostrophes, question marks, periods (written by T5 as ‘~’) and spaces (written by T5 as ‘>‘).

**Figure 2.**
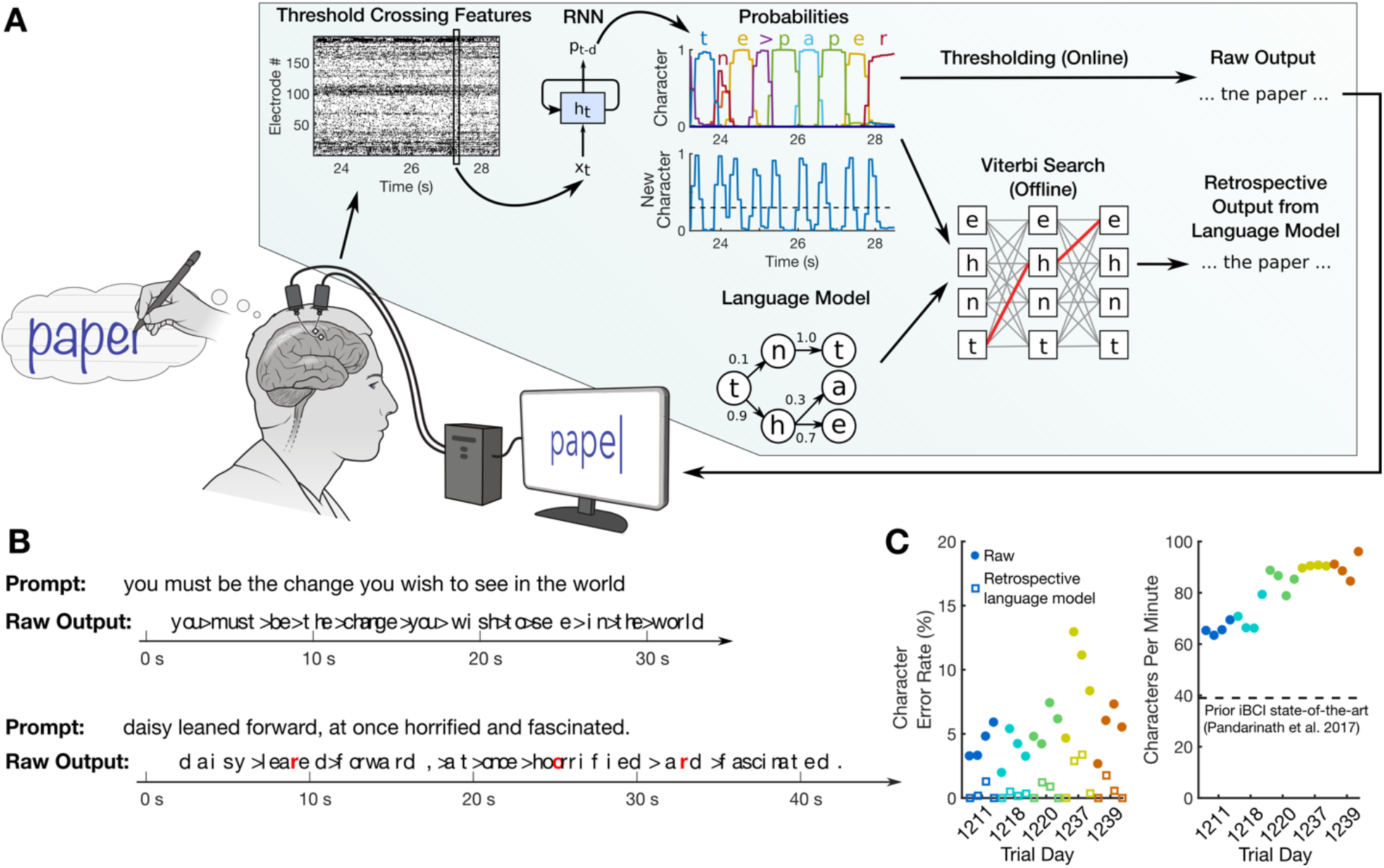
Neural decoding of attempted handwriting in real-time. **(A)** Diagram of our decoding algorithm. First, the neural activity (multiunit threshold crossings) is temporally binned (20 ms bins) and smoothed on each electrode. Then, a recurrent neural network (RNN) converts this neural population time series (*x_t_*) into a probability time series (*p_t-d_*) describing the likelihood of each character and the probability of any new character beginning. The RNN has a one second output delay (*d*) so that it has time to observe the full character before deciding its identity. Finally, the character probabilities were thresholded to produce “Raw Output” for real-time use (when the “new character” probability crossed a threshold at time *t*, the most likely character at time *t*+0.3 was emitted). In an offline retrospective analysis, the character probabilities were combined with a large-vocabulary language model to decode the most likely text that the participant wrote (we used a custom 50,000-word bigram model). **(B)** Two realtime example trials are shown, demonstrating the RNN’s ability to decode readily understandable text on sentences it was never trained on. Errors are highlighted in red and spaces are denoted with “>“. **(C)** Error rates (edit distances) and typing speeds are shown for five days, with four blocks of 7-10 sentences each (each block indicated with a single circle). The speed is more than double that of the next fastest intracortical BCI^7^.

To collect training data for the RNN, we recorded neural activity while T5 attempted to handwrite complete sentences at his own pace. A computer monitor instructed T5 which sentences to write and when to begin writing. Prior to the first day of real-time use described here, we collected a total of 242 sentences across 3 days that were combined to train the RNN (sentences were selected from the British National Corpus). After each new day of decoder evaluation, that day’s data was cumulatively added to the training dataset for the next day (yielding a total of 572 sentences by the last day). To train the RNN, we adapted neural network methods in automatic speech recognition^12–15^ to overcome two key challenges: (1) the time that each letter was written in the training data was unknown (since T5’s hand didn’t move), making it challenging to apply supervised learning techniques, and (2) the dataset was limited in size compared to typical RNN datasets, making it difficult to prevent overfitting to the training data (see Methods, Supplemental Methods, SFigs. 2-3).

We evaluated the RNN’s performance over a series of 5 days, each day containing 4 evaluation blocks of 7-10 sentences that the RNN was never trained on (thus ensuring that the RNN could not have overfit to those sentences). T5 copied each sentence from an onscreen prompt, attempting to handwrite it letter by letter, while the decoded characters appeared on the screen in real-time as they were detected by the RNN (SVideo 1, Table S2). Characters appeared after they were completed by T5 with a short delay (estimated to be between 0.4-0.7 seconds, see Methods). The decoded sentences were quite legible (Fig. 2B, “Raw Output”). Importantly, typing rates were high, plateauing at 90 characters per minute with a 5.4% character error rate (Fig. 2C, average of red circles). When a language model was used to autocorrect errors, error rates decreased considerably (Fig. 2C, open squares below filled circles; Table 1). The word error rate fell to 3.4% averaged across all days, which is comparable to state-of-the-art speech recognition systems (e.g. 4-5%^15,16^), putting it well within the range of usability. Finally, to probe the limits of possible decoding performance, we retrospectively trained a new RNN using all available sentences to process the entire sentence in a non-causal way (comparable to other BCI studies^17,18^). In this regime, accuracy was extremely high (0.17% character error rate averaged across all sentences), indicating a high potential ceiling of performance.

**Table 1.**
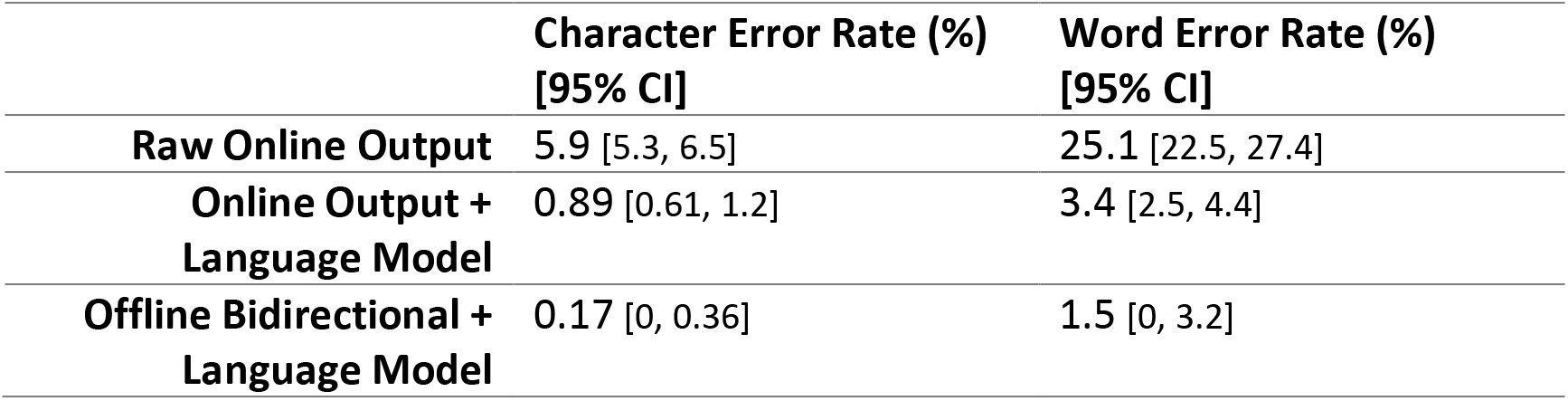
Mean character and word error rates (with 95% CIs) for the handwriting BCI across all 5 days. “Raw Online Output” is what was decoded online (in real-time). “Online Output + Language Model” was obtained by applying a language model retrospectively to what was decoded online. “Offline Bidirectional + Language Model” was obtained by retraining a bidirectional (acausal) decoder offline using all available data, in addition to applying a language model. Confidence intervals (CIs) were computed with the bootstrap percentile method (resampling over trials 10,000 times).

|  | Character Error Rate (%)<br>[95% CI] | Word Error Rate (%)<br>[95% CI] |
| --- | --- | --- |
| Raw Online Output | 5.9 [5.3, 6.5] | 25.1 [22.5, 27.4] |
| Online Output +<br>Language Model | 0.89 [0.61, 1.2] | 3.4 [2.5, 4.4] |
| Offline Bidirectional +<br>Language Model | 0.17 [0, 0.36] | 1.5 [0, 3.2] |

Next, to evaluate performance in a less restrained setting, we collected two days of data in which T5 used the BCI to freely type answers to open-ended questions (SVideo 2, Table S3). The results confirm that high performance can also be achieved when the user writes self-generated sentences as opposed to copying on-screen prompts (73.8 characters per minute with an 8.54% character error rate in real-time, 2.25% with a language model). The prior state-of-the-art for free typing in intracortical BCIs is 24.4 correct characters per minute^7^.

To our knowledge, 90 characters per minute is the highest typing rate yet reported for any type of BCI (see Discussion). For intracortical BCIs, the highest performing method has been point-and-click typing with a 2D computer cursor, peaking at 40 characters per minute^7^ (see SVideo 3 for a direct comparison). How is it that handwriting movements could be decoded more than twice as fast, with similar levels of accuracy? We theorize that point-to-point movements may be harder to distinguish from each other than handwritten letters, since letters have more variety in their spatiotemporal patterns of neural activity than do straight-line movements. To test this theory, we analyzed the spatiotemporal patterns of neural activity associated with 16 straight-line movements and 16 letters that required no lifting of the pen off the page, both performed by T5 with attempted handwriting (Fig. 3A-B).

**Figure 3.**
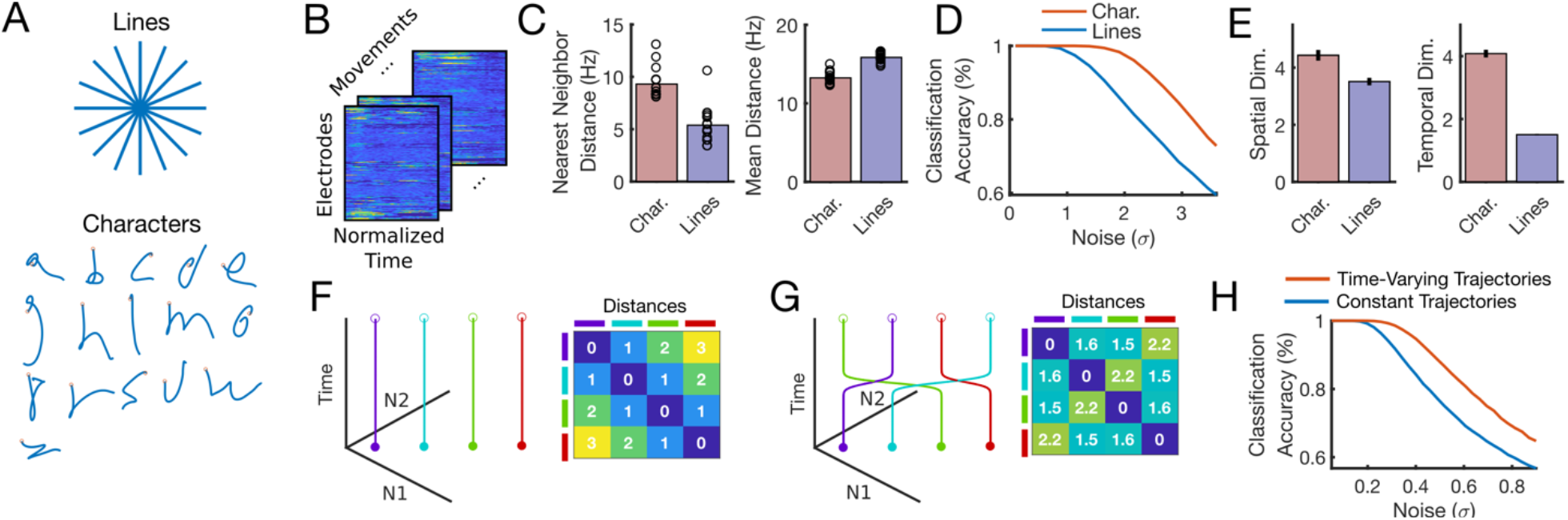
Increased temporal complexity can make movements easier to decode. (A) We analyzed the spatiotemporal patterns of neural activity corresponding to 16 handwritten characters (1 second in duration) vs. 16 handwritten straight-line movements (0.6 seconds in duration). (B) Spatiotemporal neural patterns were found by averaging over all trials for a given movement (after time-warping to align the trials in time)^11^. Neural activity was then resampled to equalize the duration of each set of movements (otherwise straight-line movements would be shorter in duration), resulting in a 192 x 100 matrix for each movement (192 electrodes and 100 time steps). (C) Pairwise Euclidean distances between neural patterns were computed for each set, revealing a larger nearest neighbor distance (but not mean distance) for characters. Each circle represents a single movement and bar heights show the mean. (D) Larger nearest neighbor distances made the characters easier to classify than straight lines. The noise is in units of standard deviations and matches the scale of the distances in C. (E) The spatial dimensionality was similar for characters and straight lines, but the temporal dimensionality was more than twice as high for characters, suggesting that more complex temporal patterning underlies the increased nearest neighbor distance and better classification performance. Error bars show the 95% CI (bootstrap percentile method). Dimensionality was defined as the participation ratio, which is approximately equal to the number of dimensions needed to explain 80% of the variance^19^. (F, G, H) A toy example can give intuition for how increased temporal dimensionality can make neural trajectories more separable. Four neural trajectories are depicted (N1 and N2 are two hypothetical neurons whose activity is constrained to a single spatial dimension, the unity diagonal). Allowing the trajectories to vary in time by adding one bend (increasing the temporal dimensionality from 1 to 2) enables larger nearest neighbor distances (G) and better classification (H).

First, we analyzed the pairwise Euclidean distances between each neural activity pattern. We found that the nearest-neighbor distances for each movement were almost twice as large for characters as compared to straight lines (72% larger), making it less likely for a decoder to confuse two nearby characters (Fig. 3C). To confirm this, we simulated the classification accuracy for each set of movements as a function of neural noise (Fig. 3D; see Methods), demonstrating that characters are easier to classify than straight lines. Note that classification accuracy begins to decrease significantly when the standard deviation of the noise is around one quarter that of the nearest neighbor distances (at this size, the clouds of noise around each point begin to intersect, resulting in decoding errors).

To gain insight into what might be responsible for the relative increase in nearest neighbor distances for characters, we examined the spatial and temporal dimensionality of the neural patterns. Spatial and temporal dimensionality were quantified using the participation ratio (PR), which quantifies approximately how many spatial or temporal axes are required to explain 80% of the variance in the neural activity patterns^19^. We found that the spatial dimensionality was similar for straight-lines and characters (Fig. 3E), but that the temporal dimensionality was more than twice as large for characters, suggesting that the increased variety of temporal patterns in letter writing drives the increased separability of each movement. To illustrate how increased temporal dimensionality can make movements more distinguishable, we constructed a toy model with four movements and two neurons whose activity is constrained to lie along a single dimension (Fig. 3F-G). Simply by allowing the trajectories to change in time (Fig. 3G), the nearest neighbor distance between the neural trajectories can be increased, resulting in an increase in classification accuracy when noise levels are large enough (Fig. 3H).

These results suggest that time-varying patterns of movement, such as handwritten letters, are fundamentally easier to decode than point-to-point movements, and can thus enable higher communication rates. This concept could be applied more generally to improve any BCI that enables discrete selection between a set of options (by associating these options with time-varying gestures as opposed to simple movements). Using the principle of maximizing the nearest neighbor distances between movements, it is possible to optimize a set of trajectories for ease of classification (as has been done previously to optimize target locations^20^). We explored doing so and designed an alphabet that is theoretically easier to classify than the Latin alphabet (SFig. 4). Our results revealed one drawback of the Latin alphabet from a neural decoding perspective: there are large clusters of redundant letters that are written similarly (most letters begin with either a downstroke or a counter-clockwise curl).

## Discussion

Commonly used BCIs for restoring communication to people who can’t move or speak are either flashing EEG spellers^21–26^ or 2D point-and-click computer cursor-based BCIs for selecting letters on a virtual keyboard^27,6,7^. EEG spellers that use visually evoked potentials have achieved speeds of 60 characters per minute^24^, but have important usability limitations, as they tie up the eyes, are not typically self-paced, and require panels of flashing lights on a screen that take up space and may be fatiguing. Intracortical BCIs based on 2D cursor movements give the user more freedom to look around and set their own pace of communication, but have yet to exceed 40 correct characters per minute in people^7^. Recently, speech-decoding BCIs have shown exciting promise for restoring rapid communication (e.g.^28,17,18^), but their accuracies and vocabularies are currently too limited for general-purpose use.

Here, we introduced a novel approach for communication BCIs – decoding a rapid, dexterous motor behavior in a person with paralysis – that, to our knowledge, set a new standard for communication rate at 90 characters per minute. We demonstrated a real-time system that is general (the user can express any sentence), easy to use (entirely self-paced and the eyes are free to move), and accurate enough to be useful in the real-world (94.5% raw accuracy and >99% with a language model capable of running in real-time). We anticipate that a handwriting decoder could be combined with a high-performance point-and-click decoder^7^ to enable both rapid typing and general-purpose use of computer applications. One unique advantage of our handwriting BCI is that, in theory, it does not require vision (since no feedback of the imagined pen trajectory is given to the participant, and letters appear only after they are completed). To our knowledge, it is the first high-speed BCI that has the potential to work in people with visual impairments.

To achieve high performance, we developed new decoding methods to overcome two key challenges: (1) lack of observable behavior during long sequences of self-paced training data (our participant’s hand never moved), and (2) limited amounts of training data. Our techniques adapt neural network methods from automatic speech recognition^12–15^ to work effectively with neural activity in data-limited regimes. These methods could be useful more generally for neurally decoding any sequential behavior that cannot be observed directly (for example, decoding speech from someone who can no longer speak).

Finally, it is important to recognize that our system is a proof-of-concept that a high-performance handwriting BCI is possible; it is not yet a complete, clinically viable system. Current limitations include a reduced character set (e.g. no capital letters), inability to delete or edit text, and a relatively long calibration process (although see SFig. 5, where we show retrospectively that good performance can still be achieved with less calibration time). To facilitate further investigation and refinement, we plan to publicly release the core dataset used here to train and evaluate our handwriting decoder upon publication in a peer-reviewed journal. This unique dataset contains >40k characters over 10 days and provides a rich testbed for developing new decoding approaches.

## Acknowledgements

We thank participant T5 and his caregivers for their dedicated contributions to this research, and N. Lam, E. Siauciunas, and B. Davis for administrative support. This work was supported by the Howard Hughes Medical Institute (F.R.W. and D.T.A.), Office of Research and Development, Rehab. R&D Service, Department of Veterans Affairs (B6453R, N2864C), NIH-NIDCD R01DC014034, NIH-NINDS UH2NS095548, NIH-NINDS U01NS098968, The Executive Committee on Research (ECOR) of Massachusetts General Hospital MGH Deane Institute for Integrated Research on Atrial Fibrillation and Stroke (L.R.H.); NIDCD R01-DC014034, NIDCD U01-DC017844, NINDS UH2-NS095548, NINDS UO1-NS098968, Larry and Pamela Garlick, Samuel and Betsy Reeves, Wu Tsai Neurosciences Institute at Stanford (J.M.H and K.V.S); Simons Foundation Collaboration on the Global Brain 543045 and Howard Hughes Medical Institute Investigator (K.V.S). The funders had no role in study design, data collection and interpretation, or the decision to submit the work for publication.

## Author Contributions

F.R.W. conceived the study, designed the experiments, built the real-time decoder, analyzed the data, and wrote the manuscript. F.R.W. and D.T.A. collected the data. L.R.H. is the sponsor-investigator of the multi-site clinical trial. J.M.H. planned and performed T5’s array placement surgery and was responsible for his ongoing clinical care. J.M.H. and K.V.S. supervised and guided the study. All authors reviewed and edited the manuscript.

## Declaration of Interests

The MGH Translational Research Center has a clinical research support agreement with Neuralink, Paradromics, and Synchron, for which L.R.H. provides consultative input. JMH is a consultant for Neuralink Corp and Proteus Biomedical, and serves on the Medical Advisory Board of Enspire DBS. KVS consults for Neuralink Corp. and CTRL-Labs Inc. (part of Facebook Reality Labs) and is on the scientific advisory boards of MIND-X Inc., Inscopix Inc., and Heal Inc. All other authors have no competing interests.

## Methods

### Study Participant

This study includes data from one participant (identified as T5) who gave informed consent and was enrolled in the BrainGate2 Neural Interface System clinical trial (ClinicalTrials.gov Identifier: NCT00912041, registered June 3, 2009). This pilot clinical trial was approved under an Investigational Device Exemption (IDE) by the US Food and Drug Administration (Investigational Device Exemption #G090003). Permission was also granted by the Institutional Review Boards of Stanford University (protocol #20804). All research was performed in accordance with relevant guidelines/regulations. The BrainGate2 trial’s purpose is to collect preliminary safety information and demonstrate feasibility that an intracortical BCI can be used by people with tetraplegia for communication and control of external devices; the present manuscript results from analysis and decoding of neural activity recorded during the participants’ engagement in research that is enabled by the clinical trial but does not report clinical trial outcomes.

T5 is a right-handed man, 65 years old at the time of data collection, with a C4 AIS C spinal cord injury that occurred approximately 9 years prior to study enrollment. In August 2016, two 96 electrode intracortical arrays (Neuroport arrays with 1.5-mm electrode length, Blackrock Microsystems, Salt Lake City, UT) were placed in the hand “knob” area of T5’s left-hemisphere (dominant) precentral gyrus. Data are reported from post-implant days 994 to 1246 (Table S1). T5 retained full movement of the head and face and the ability to shrug his shoulders. Below the injury, T5 retained some very limited voluntary motion of the arms and legs that was largely restricted to the left elbow; however, some micromotions of the right hand were visible during attempted handwriting (see^10^ for neurologic exam results and SVideo 4 for hand micromotions). Array placement locations registered to MRI-derived brain anatomy are shown in^10^.

### Neural Signal Processing

Neural signals were recorded from the microelectrode arrays using the NeuroPort™ system (Blackrock Microsystems; more details are described in^6,7,29^). Neural signals were analog filtered from 0.3 Hz to 7.5 kHz and digitized at 30 kHz (250 nV resolution). Next, a common average reference filter was applied that subtracted the average signal across the array from every electrode in order to reduce common mode noise. Finally, a digital bandpass filter from 250 to 3000 Hz was applied to each electrode before threshold crossing detection.

We used multiunit threshold crossing rates as neural features for analysis and neural decoding (as opposed to spike-sorted single units). Using multiunit threshold crossings allowed us to leverage information from more electrodes, since many electrodes recorded activity from multiple neurons that could not be precisely spike-sorted into single units. Recent results indicate that neural population structure can be accurately estimated from threshold crossing rates alone^30^, and that neural decoding performance is similar to using sorted units^31^. For threshold crossing detection, we used a −3.5 x RMS threshold applied to each electrode, where RMS is the electrode-specific root mean square of the voltage time series recorded on that electrode. Threshold crossing times were “binned” into 10 ms bins (for analysis) or 20 ms bins (for decoding) to estimate the threshold crossing rate in each bin (the estimated rate was equal to the number of threshold crossings divided by the bin width).

### Session Structure and Tasks

Neural data were recorded in 3-5 hour “sessions” on scheduled days, which typically occurred 2-3 times per week. During the sessions, T5 sat upright in a wheelchair with his hand resting on his lap. A computer monitor placed in front of T5 indicated which sentence (or single character) to write and when. Data were collected in a series of 5-10 minute “blocks” consisting of an uninterrupted series of trials. In between these blocks, T5 was encouraged to rest as needed. The software for running the experimental tasks, recording data, and implementing the real-time decoding system was developed using MATLAB and Simulink (MathWorks, Natick, MA).

#### Instructed Delay Paradigm

The data were collected across 11 sessions (Table S1 details the data collected in each session). All tasks employed an instructed delay paradigm. For the single character writing task shown in Figure 1A, the delay period duration was drawn from an exponential distribution (mean of 1.5 s); values that fell outside of the range of 2.0 – 3.0 s were re-drawn. After the delay period, the text prompt changed to “Go” and the red square (stop cue) turned green for 1 second, cueing T5 to begin attempting to write.

During sentence writing blocks, the delay period always lasted 5 seconds. During this delay period the sentence to be written was displayed, providing T5 time to read the sentence. The red stop cue then turned green, and the sentence remained displayed on the screen while T5 attempted to handwrite it letter by letter. When T5 finished writing the sentence, he turned his head to the right, which our system detected and automatically triggered the next sentence. Head position was tracked optically with the OptiTrack V120:Trio bar (Corvallis, OR) containing three infrared cameras that tracked the position of optical markers worn on a head band.

#### Decoder Evaluation

Sentence-writing days where real-time decoding was tested (sessions 3-11) had the following structure (illustrated in SFig. 2A). First, we collected interleaved blocks of single character writing (2 blocks, 5 repetitions of each character per block) and sentence writing (5 blocks, 10 sentences per block); no decoder was active during these blocks. Then, we trained the decoder using these blocks of data (combined with data from all past sessions). Finally, we collected evaluation blocks where T5 used the decoder to copy sentences (sessions 3-9, 4 blocks per session) or freely answer questions (sessions 10-11, 3 blocks per session). Note that the reported data in Figure 2 are from sessions 5-9, since sessions 3-4 were pilot sessions devoted to exploring different decoding approaches. Session 1 was the only session where sentences were written but no real-time decoding was performed; in session 1, we collected 102 sentences plus 27 repetitions of each character written individually. These data were used for Figure 1 and to initialize the decoder for session 3.

For two of the four evaluation blocks in the copy typing sessions, we used the 7 sentences employed in^7^ for a direct comparison to this prior state-of-the-art point-and-click typing BCI (SVideo 3). The other two evaluation blocks contained 10 unique sentences selected from the British National Corpus (BNC)^32^. Sentences were chosen from the BNC (using the Sketch Engine tool) by first randomly selecting words from a list of the top 2,000 most common words in the BNC. Then, for each randomly chosen word, the BNC was searched for example sentences containing that word; we hand-selected examples of reasonable length (no more than 120 characters) and whose meaning was not too confusing out of context, so as not to be distracting to T5. The end result was a diverse sample of sentences from many different contexts (spoken English, fiction, non-fiction, news, etc.). Finally, we added 5 pangrams (sentences containing all 26 letters) to each session’s training data that did not appear in the BNC, in order to increase the frequency of rare letters.

To prevent the decoder from artificially increasing its performance by overfitting to specific sentences, our decoder was never evaluated on a sentence that it had been previously trained on, and every sentence was unique (except for the “direct comparison” blocks containing sentences taken from^7^). We excluded these direct comparison blocks from the RNN’s training dataset, so that it could not overfit to these repeated sentences.

The two free-typing sessions (sessions 10-11) used 8 blocks of sentence-writing data for decoder training (instead of 5) and used a different set of sentences to add more realistic variability to the training data. For 3 of the 8 sentence writing blocks, we randomly added hash mark characters (#) throughout the BNC sentences, which signaled T5 to take an artificial pause from writing. For the other 5 blocks, we used short 2-4 word phrases instead of complete sentences and asked T5 to write from memory (instead of copying what was on the screen). To enforce writing from memory, we removed the phrase from the screen during the “Go” period.

### Pen Trajectory Visualization

To make Figure 1, threshold crossing rates were first binned into 10 ms bins and smoothed by convolving with a gaussian kernel (sd = 30 ms) to remove high frequency noise. The smoothed rates were then compiled into a matrix of dimension N x TC, where N is the number of microelectrodes (192), T is the number of 10 ms time bins (200), and C is the number of characters (31). Each row contains the trial-averaged response of a single electrode to each character, in a time window from −500 ms to 1500 ms around the go cue (the 31 trial-averaged responses were concatenated together into a single vector). Principal components analysis was then applied to this matrix to find the top 3 PCs which were used to visualize the raw activity (Figure 1B).

Next, we used time-warped PCA (https://github.com/ganguli-lab/twpca)^11,33^ to find continuous, regularized time-warping functions that align the trials within a single movement condition together. We verified that these warping functions appeared close to the identity line, smoothly bending away from it after the go cue in order to account for variations in writing speed from trial to trial (as can be seen in the example shown in Figure 1B-C). We used the following time-warping parameters: 5 components, 0.001 scale warping regularization (L1), and 1.0 scale time regularization (L2).

Finally, we trained a linear decoder to readout pen tip velocity from the neural activity. The decoder computed velocity as follows:

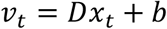

Here, *v_t_* is a 2 x 1 pen tip velocity vector containing X and Y velocity at time t, *D* is a 2 x 192 decoding matrix, *x_t_* is a 192 x 1 vector of binned threshold crossing rates, and *b* is a 2 x 1 offset term. Importantly, each character was held-out when decoding in a leave-one-out fashion. That is, the pen tip velocities for any given character were obtained using a decoder that was trained on all *other* characters, preventing the decoder from trivially overfitting and reproducing the templates used to train it.

To train the decoder, we used hand-made templates that describe each character’s pen trajectory. The character templates were made by drawing each character with a computer mouse in the same way as T5 described writing the character; these templates then defined the target velocity vector for the decoder on each time step of each trial. We used ordinary least squares regression to train the decoder to minimize the error between the template velocities and the decoded velocities (see Supplemental Methods for more details).

### t-SNE Visualization and k-nearest neighbor classifier

For Figure 1E, we used t-distributed stochastic neighbor embedding (t-SNE)^34^ to nonlinearly reduce the dimensionality of trials of neural activity for visualization (perplexity=50). Before applying t-SNE, we smoothed the neural activity and reduced its dimensionality to 15 with PCA (using the methods described in the above section). Each trial of neural activity was thus represented by a 200 x 15 matrix (200 time bins by 15 dimensions). We applied t-SNE to these matrices using the following “time-warp” distance function:

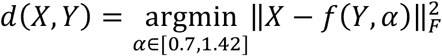

Here, α is a time-warp factor, *f* is a warping function, and *X* and *Y* are trials of neural activity. The function *f* time-warps a trial of neural activity by resampling with linear interpolation by a factor of α. After warping, only the first *N* shared time bins between *X* and *Y* are used to compute the distance. The warping function serves to account for differences in writing speed across trials, so that the same pattern of neural activity occurring at a different speed is considered nearby.

To make the t-SNE plot clearer, we removed a small number of outliers from each class, which resulted in removing 3% of data points that were likely caused by lapsed attention by T5. Outliers were defined as having a mean within-class pairwise distance that was greater than 4 median absolute deviations from average.

We also used the time-warp distance function to perform k-nearest neighbor classification (k=10), which resulted in 94.1% accuracy (compared to 88.8% accuracy with a Euclidean distance function).

### Recurrent Neural Network Decoder

We used a two layer, gated recurrent unit RNN^35^ to convert T5’s neural activity into a time series of character probabilities (see SFig. 1 for a diagram). We found that a recurrent neural network decoder strongly outperformed using a simple hidden Markov model decoder (Table S4).

As a pre-processing step, threshold crossing rates were binned in 20 ms time steps, z-scored, causally smoothed by convolving with a gaussian kernel (sd = 40 ms) that was delayed by 100 ms, and concatenated into a 192 x 1 vector *x_t_*. We used the following variant of the gated recurrent unit RNN that is implemented by the cuDNN library^36^:

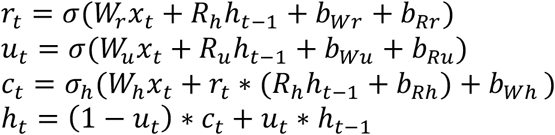

Here, *σ* is the logistic sigmoid function, *σ_h_* is the hyperbolic tangent, *x_t_* is the input vector at time step *t*, *h_t_* is the hidden state vector, *r_t_* is the reset gate vector, *u_t_* is the update gate vector, *c_t_* is the candidate hidden state vector, *W, R* and *b* are parameter matrices and vectors, and * denotes the element-wise multiplication.

We used a two-layer RNN architecture (where the hidden state of the first layer was fed as input to the second layer). Importantly, the RNN was trained with an output delay. That is, the RNN was trained to predict the character probabilities from 1 second in the past; this was necessary to ensure that the RNN had enough time to process the entire character before deciding on its identity. The output probabilities were computed from the hidden state of the second layer as follows:

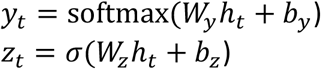

Here, *σ* is the logistic sigmoid function, *h_t_* is the hidden state of the second layer, *W* and *b* are parameter matrices and vectors, *y_t_* is a vector of character probabilities (one entry for each character), and *z_t_* is a scalar probability that represents the probability of any new character beginning at that time step. During real-time operation, we thresholded *z_t_* (threshold = 0.3) to decide when to emit a new character. Whenever *z_t_* crossed the threshold, we emitted the most probable character in *y_t_* 300 ms later.

We updated the second layer of our RNN decoder at a slower frequency than the first layer (every five 20 ms time steps instead of every single time step). We found that this increased the speed and reliability of training, making it easier to hold information in memory for the length of the output delay (e.g., for a 1 second delay, the slower frequency means that the top layer must hold information in memory for only 10 steps as opposed to 50).

### RNN Training

See SFig. 2B for a diagram of the RNN training flow and Supplemental Methods for a detailed protocol. Here, we give an overview of the main steps and algorithms used. Our methods are an adaptation of neural network methods used in automatic speech recognition^12,37,14,15^, with key changes to achieve high performance on neural activity in a highly data-limited regime (1-10 hours of data, compared to 1-10k hours, e.g.^38–40^).

#### Data Labeling

A major challenge that had to be overcome for training decoders with our data is that we don’t know what character T5 was writing at any given moment in time in the training data. There are two major approaches used to solve this problem in automatic speech recognition: forced-alignment labeling with hidden Markov Models (HMMs)^41,12,15^, or unsupervised inference with connectionist temporal classification^42^ (and other similar cost functions, e.g.^43^). We found that forced-alignment worked better with our data, potentially because of the relatively small dataset size; it also enabled data augmentation via synthetic sentence generation (see below). In the forced-alignment method, HMMs are used to infer what character is being written at each time step, fusing knowledge of the sequence of characters that were supposed to be written with the neural activity recorded. These character labels can then be used to construct target probabilities that the RNN is trained to reproduce in a supervised manner.

To construct the data-labeling HMMs, we first processed the single character data to convert it into trial-averaged spatiotemporal “templates” of the neural activity patterns associated with each character. Next, these templates were used to define the emission probabilities of the HMMs, and the states and transition probabilities were set to express an orderly march through the sequence of characters in each sentence (SFig. 2C). We then used the Viterbi algorithm to find the most probable start time of each character given the observed neural activity. The start times of each character could then be used to construct target time series of character probabilities for the RNN to reproduce.

The vector of target character probabilities (denoted as *y_t_* above) was constructed by setting the probability at each time step to be a one-hot representation of the most recently started character. The scalar character start probability (denoted as *z_t_* above) was set to be equal to 1 for a 200 ms window after each character began, and was otherwise equal to 0. The character start probability allows the decoder to distinguish repeated characters from single characters (e.g., “oo” vs. “o”).

One advantage of this strategy for representing the RNN output is that uncertainty about whether pauses are occurring between characters should not degrade performance, since the labeling routine only needs to identify when each character begins (not when it ends). Note that this representation causes the RNN to output a “sample-and-hold”-type signal, where it will continue to output the most recently started character until the next one begins.

#### Supervised Training

Once the data were labeled, we used those labels to cut out snippets of each character from the data. These snippets were then re-assembled into artificial sentences, which were added to the training data to augment it and prevent overfitting (SFig. 2E). This data augmentation step was critical for achieving high performance (SFig. 3A). Finally, with the labeled and augmented dataset, standard supervised training techniques were then employed to train the RNN. We used TensorFlow v1.15^44^ to train the RNN with gradient descent (using Adam^45^). To train the RNN to account for changes in the means of neural features which naturally occur over time^6,46^, we added artificial perturbations to the feature means (similar to^47^). This step was also essential to achieving high-performance (SFig. 3B).

On each new day, we re-trained the RNN to incorporate that new day’s data before doing real-time performance evaluation. The new data were combined with all previous days’ data into one large dataset while training. To account for differences in neural activity across days^6,48^, we separately transformed each days’ neural activity with a linear transformation that was simultaneously optimized with the other RNN parameters. Including multiple days of data, and fitting separate input layers for each day, substantially improved performance (SFig. 3C-D).

### Language Model

In a retrospective analysis, we used a custom, large vocabulary language model to autocorrect errors made by the decoder. Here, we give an overview of the major steps involved (see Supplemental Methods for details). The language model had two stages: (1) a 50,000-word bigram model that first processes the neural decoder’s output to generate a set of candidate sentences, and (2) a neural network to rescore these candidate sentences (OpenAI’s GPT-2, 1558M parameter version; https://github.com/openai/gpt-2)^49^. This two-step strategy is typical in speech recognition^15^ and plays to the strengths of both types of models. Although the rescoring step improved performance, we found that performance was strong with the bigram model alone (1.48% character error rate with the bigram model alone, 0.89% with rescoring).

The bigram model was created with Kaldi^50^ using samples of text provided by OpenAI (250k samples from WebText, https://github.com/openai/gpt-2-output-dataset). These samples were first processed to make all text lower case and to remove all punctuation that was not part of our limited character set (consisting only of periods, question marks, commas, apostrophes, and spaces). Then, we used the Kaldi toolkit to construct a bigram language model, using the 50,000 most common words appearing the WebText sample, in the form of a finite-state transducer which could be used to translate the RNN probabilities into candidate sentences^51^.

### Performance Metrics

Character error rate was defined as the edit distance between the decoded sentence and the prompt (i.e., the number of insertions, deletions or substitutions required to make the strings of characters match exactly). Similarly, word error rate was the edit distance defined over sequences of “words” (strings of characters separated by spaces; punctuation was included as part of the word it appeared next to). For the free typing sessions, the intended sentence was determined by discussing with the participant his intended meaning.

Characters per minute was defined as 60(E-S)/N, where N was the number of characters in the target sentence, E was the end time and S was the start time (in seconds). For copy typing, S was the time of the go cue and E was the time of the last decoded character. Rarely, T5 had lapses of attention and did not respond to the go cue until many seconds later; we thus capped his reaction time (the time between the go cue and the first decoded character) to no more than 2 seconds. For the free typing sessions, we defined S as the time of the first decoded character (instead of the go cue), since T5 often took substantial time after the go cue to formulate his response to the prompt.

We removed a small number of trials with incomplete data (5%); this occurred during the copy typing sessions when T5 accidentally triggered the next sentence by moving his head too far to the right before he had finished typing the sentence (our system triggered the next sentence upon detection of a rightward head turn). During free typing, we removed one sentence where T5 could not think of a response and wanted to skip the question.

To estimate the able-bodied smartphone typing rate of people in T5’s age group (115 characters per minute, as mentioned in the Summary), we used the publicly available data from^8^. We took the median over all participants greater than 60 years old (T5 was 65 at the time of data collection).

### Estimating the Time Between Character Completion and On-Screen Appearance

We estimate that characters were emitted between 0.4-0.7 seconds after T5 completed them. Our logic is as follows. When T5 begins attempting to write a new character, our decoder takes 1 second to recognize this (as it is trained with 1 second delay). Thus, it will emit a new character start signal (*z_t_*) approximately 1 second after the character begins. Adding an additional 0.1 seconds for the causal gaussian smoothing delay and 0.3 seconds to emit the character after *z_t_* crosses threshold yields a 1.4 second delay between when the character is started and when it appears on the screen. This means the character will appear on the screen ~ 1.4 – X seconds after T5 completes it, where X is the time taken to write the letter. For a 90 characters per minute typing rate, characters take on average 60/90 = 0.66 seconds to write, thus yielding a 1.4 – 0.66 = 0.74 second delay. For slower typing rates (e.g., 60 characters per minute), the delay is shorter (~0.4 seconds).

### Temporal Complexity Increases Decodability

#### Pairwise Neural Distances

The characters dataset analyzed in Figure 3 is the same as that shown in Figure 1 (session 1). The straight-lines dataset was collected on a separate session (session 2; see Table S1), where T5 attempted to handwrite straight-line strokes in an instructed delay paradigm identical to what was used for writing single characters (except instead of a text cue, a line appeared on the screen to indicate the direction of the stroke).

To compute the pairwise distances reported in Figure 3C, the threshold crossing rates were first binned into 10 ms bins and smoothed by convolving with a Gaussian kernel (30 sd). Then, neural activity within a 0 to 1000 ms window after the go cue (for characters) or 0 to 600 ms (for lines) was time-aligned across trials using the time-warping methods described above (for Figure 1). These time windows were chosen by visual inspection of when the neural activity stopped modulating. The time-aligned data were then trial-averaged and re-sampled to 100 time points (using linear interpolation) to generate a set of mean spatiotemporal neural activity matrices (of dimension 192 electrodes x 100 time steps).

Pairwise distances were defined as the Euclidean norm (square root of the sum of squared entries) of the difference matrix obtained by subtracting one spatiotemporal neural matrix from another. Pairwise distances were estimated using cross-validation, according to the methods in^10^ (https://github.com/fwillett/cvVectorStats); without cross-validation, noise would inflate the distances and make them all appear larger than they are. Pairwise distances for simulated data (Figure 3F-G and SFig. 4) were computed without cross-validation (because there was no estimation noise).

We normalized the pairwise distances reported in Figure 3C by the number of time steps included in the analysis and the number of electrodes by dividing by 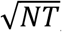, where N is the number of electrodes (192) and T is the number of time steps (100). This makes the distances roughly invariant to the number of time steps and electrodes; for example, if each electrode fires at 150 Hz for condition A and 50 Hz for condition B, then the distance between B and A is 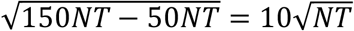.

#### Neural and Temporal Dimensionality

Dimensionality, as computed in Figure 3E, was estimated using the “participation ratio”^19^, which is a continuous metric that quantifies how evenly-sized the eigenvalues of the covariance matrix are. It is roughly equivalent to the number of dimensions needed to explain 80% of the variance in the data.

To compute the neural dimensionality, the smoothed, time-warped, and trial-averaged neural activity was arranged into a matrix X of dimensionality N x TC, where N is the number of electrodes (192), T is the number of time steps (100), and C is the number of movement conditions (31). Each row is the trial-averaged response of a single electrode to each movement condition concatenated together. The eigenvalues *u_i_* of the covariance matrix XX^T^ were then used to compute the participation ratio:

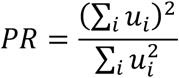

Similarly, to compute the temporal dimensionality, the neural activity was arranged into a matrix X of dimensionality T x NC, where each row contains the trial-averaged response of all neurons across all conditions concatenated together for a single time step (and each column is a neural response for a single condition). Roughly, the temporal dimensionality quantifies how many T-dimensional neural response basis vectors are needed to explain 80% of the variance of all of the neural responses (PSTHs).

To reduce bias, we used cross-validation to estimate the covariance matrix (otherwise, the presence of estimation noise would artificially inflate the dimensionality). Specifically, we split the trials into two folds, and computed the X matrix separately for each fold (yielding X1 and X2). The covariance matrix was then estimated as X_1_X_2_^T^. To compute confidence intervals, we used the jackknife method (see our prior work^10^ https://github.com/fwillett/cvVectorStats).

#### Simulated Classification Accuracy

To simulate classification accuracy for the lines and characters as a function of neural noise (Figure 3D), we used the cross-validated pairwise distances between all conditions. This ensures that accuracy is not inflated by overestimating the distances between conditions. We used classic multidimensional scaling to find a set of low-dimensional points that have the same pairwise distances; these are low-dimensional representations of the neural activity patterns associated with each movement class. Then, we simulated a trial of classification with these points by first picking a point at random (the true class), and then adding Gaussian white noise to this point in the low-dimensional space (to generate the observation). Classification was correct if the observation lay closest to the true point. This simulates a simple classifier that chooses which class an observation belongs to by choosing the class with the nearest mean (this corresponds to a maximum likelihood classifier in the case of spherical Gaussian noise).

When dealing with simulated data (Figure 3F-G or SFig. 4), no multidimensional scaling was needed since no estimation noise was present. Thus, we performed the simulated classification on the true neural patterns themselves.

## References

1. Hochberg, L. R. et al. Reach and grasp by people with tetraplegia using a neurally controlled robotic arm. Nature 485, 372–375 (2012).

2. Collinger, J. L. et al. High-performance neuroprosthetic control by an individual with tetraplegia. The Lancet 381, 557–564 (2013).

3. Aflalo, T. et al. Decoding motor imagery from the posterior parietal cortex of a tetraplegic human. Science 348, 906–910 (2015).

4. Bouton, C. E. et al. Restoring cortical control of functional movement in a human with quadriplegia. Nature 533, 247–250 (2016).

5. Ajiboye, A. B. et al. Restoration of reaching and grasping movements through brain-controlled muscle stimulation in a person with tetraplegia: a proof-of-concept demonstration. The Lancet 389, 1821–1830 (2017).

6. Jarosiewicz, B. et al. Virtual typing by people with tetraplegia using a self-calibrating intracortical brain-computer interface. Science Translational Medicine 7, 313ra179–313ra179 (2015).

7. Pandarinath, C. et al. High performance communication by people with paralysis using an intracortical brain-computer interface. eLife 6, e18554 (2017).

8. Palin, K., Feit, A. M., Kim, S., Kristensson, P. O. & Oulasvirta, A. How do People Type on Mobile Devices? Observations from a Study with 37,000 Volunteers. in Proceedings of the 21st International Conference on Human-Computer Interaction with Mobile Devices and Services 1–12 (Association for Computing Machinery, 2019). doi:10.1145/3338286.3340120.

9. Yousry, T. A. et al. Localization of the motor hand area to a knob on the precentral gyrus. A new landmark. Brain 120, 141–157 (1997).

10. Willett, F. R. et al. Hand Knob Area of Premotor Cortex Represents the Whole Body in a Compositional Way. Cell (2020) doi:10.1016/j.cell.2020.02.043.

11. Williams, A. H. et al. Discovering Precise Temporal Patterns in Large-Scale Neural Recordings through Robust and Interpretable Time Warping. Neuron 105, 246–259.e8 (2020).

12. Hinton, G. et al. Deep Neural Networks for Acoustic Modeling in Speech Recognition: The Shared Views of Four Research Groups. IEEE Signal Processing Magazine 29, 82–97 (2012).

13. Graves, A., Mohamed, A. & Hinton, G. Speech recognition with deep recurrent neural networks. in 2013 IEEE International Conference on Acoustics, Speech and Signal Processing 6645–6649 (2013). doi:10.1109/ICASSP.2013.6638947.

14. Zeyer, A., Doetsch, P., Voigtlaender, P., Schlüter, R. & Ney, H. A comprehensive study of deep bidirectional LSTM RNNS for acoustic modeling in speech recognition. in 2017 IEEE International Conference on Acoustics, Speech and Signal Processing (ICASSP) 2462–2466 (2017). doi:10.1109/ICASSP.2017.7952599.

15. Xiong, W. et al. The Microsoft 2017 Conversational Speech Recognition System. arXiv:1708.06073 [cs] (2017).

16. He, Y. et al. Streaming End-to-end Speech Recognition for Mobile Devices. in ICASSP 2019 - 2019 IEEE International Conference on Acoustics, Speech and Signal Processing (ICASSP) 6381–6385 (2019). doi:10.1109/ICASSP.2019.8682336.

17. Anumanchipalli, G. K., Chartier, J. & Chang, E. F. Speech synthesis from neural decoding of spoken sentences. Nature 568, 493–498 (2019).

18. Makin, J. G., Moses, D. A. & Chang, E. F. Machine translation of cortical activity to text with an encoder-decoder framework. Nature Neuroscience 1–8 (2020) doi:10.1038/s41593-020-0608-8.

19. Gao, P. et al. A theory of multineuronal dimensionality, dynamics and measurement. bioRxiv 214262 (2017) doi:10.1101/214262.

20. Cunningham, J. P., Yu, B. M., Gilja, V., Ryu, S. I. & Shenoy, K. V. Toward optimal target placement for neural prosthetic devices. J. Neurophysiol. 100, 3445–3457 (2008).

21. Vansteensel, M. J. et al. Fully Implanted Brain–Computer Interface in a Locked-In Patient with ALS. New England Journal of Medicine 375, 2060–2066 (2016).

22. Nijboer, F. et al. A P300-based brain–computer interface for people with amyotrophic lateral sclerosis. Clinical Neurophysiology 119, 1909–1916 (2008).

23. Townsend, G. et al. A novel P300-based brain–computer interface stimulus presentation paradigm: Moving beyond rows and columns. Clinical Neurophysiology 121, 1109–1120 (2010).

24. Chen, X. et al. High-speed spelling with a noninvasive brain–computer interface. Proc Natl Acad Sci U S A 112, E6058–E6067 (2015).

25. McCane, L. M. et al. P300-based brain-computer interface (BCI) event-related potentials (ERPs): People with amyotrophic lateral sclerosis (ALS) vs. age-matched controls. Clinical Neurophysiology 126, 2124–2131 (2015).

26. Wolpaw, J. R. et al. Independent home use of a brain-computer interface by people with amyotrophic lateral sclerosis. Neurology 91, e258–e267 (2018).

27. Bacher, D. et al. Neural Point-and-Click Communication by a Person With Incomplete Locked-In Syndrome. Neurorehabil Neural Repair 29, 462–471 (2015).

28. Mugler, E. M. et al. Direct classification of all American English phonemes using signals from functional speech motor cortex. J. Neural Eng. 11, 035015 (2014).

## Methods References

29. Hochberg, L. R. et al. Neuronal ensemble control of prosthetic devices by a human with tetraplegia. Nature 442, 164–171 (2006).

30. Trautmann, E. M. et al. Accurate estimation of neural population dynamics without spike sorting. Neuron (2019).

31. Christie, B. P. et al. Comparison of spike sorting and thresholding of voltage waveforms for intracortical brain–machine interface performance. J. Neural Eng. 12, 016009 (2014).

32. The British National Corpus, version 3 (BNC XML Edition). (2007).

33. Poole, B. et al. Time-warped PCA: simultaneous alignment and dimensionality reduction of neural data. Cosyne Abstracts (2017).

34. Maaten, L. van der & Hinton, G. Visualizing Data using t-SNE. Journal of Machine Learning Research 9, 2579–2605 (2008).

35. Cho, K. et al. Learning Phrase Representations using RNN Encoder-Decoder for Statistical Machine Translation. arXiv:1406.1078 [cs, stat] (2014).

36. Chetlur, S. et al. cuDNN: Efficient Primitives for Deep Learning. arXiv:1410.0759 [cs] (2014).

37. Graves, A., Mohamed, A. & Hinton, G. Speech recognition with deep recurrent neural networks. in 2013 IEEE International Conference on Acoustics, Speech and Signal Processing 6645–6649 (2013). doi:10.1109/ICASSP.2013.6638947.

38. Cieri, C., Miller, D. & Walker, K. The Fisher Corpus: a Resource for the Next Generations of Speech-to-Text. in Proceedings of the Fourth International Conference on Language Resources and Evaluation (LREC’04) (European Language Resources Association (ELRA), 2004).

39. Panayotov, V., Chen, G., Povey, D. & Khudanpur, S. Librispeech: An ASR corpus based on public domain audio books. in 2015 IEEE International Conference on Acoustics, Speech and Signal Processing (ICASSP) 5206–5210 (2015). doi:10.1109/ICASSP.2015.7178964.

40. He, Y. et al. Streaming End-to-end Speech Recognition for Mobile Devices. in ICASSP 2019 - 2019 IEEE International Conference on Acoustics, Speech and Signal Processing (ICASSP) 6381–6385 (2019). doi:10.1109/ICASSP.2019.8682336.

41. Young, S. J. et al. The HTK book version 3.4. (2006).

42. Graves, A., Fernández, S., Gomez, F. & Schmidhuber, J. Connectionist temporal classification: labelling unsegmented sequence data with recurrent neural networks. in Proceedings of the 23rd international conference on Machine learning 369–376 (Association for Computing Machinery, 2006). doi:10.1145/1143844.1143891.

43. Collobert, R., Puhrsch, C. & Synnaeve, G. Wav2Letter: an End-to-End ConvNet-based Speech Recognition System. arXiv:1609.03193 [cs] (2016).

44. Abadi, M. et al. TensorFlow: Large-Scale Machine Learning on Heterogeneous Distributed Systems. arXiv:1603.04467[cs] (2016).

45. Kingma, D. P. & Ba, J. Adam: A Method for Stochastic Optimization. arXiv:1412.6980[cs] (2017).

46. Downey, J. E., Schwed, N., Chase, S. M., Schwartz, A. B. & Collinger, J. L. Intracortical recording stability in human brain-computer interface users. J. Neural Eng. 15, 046016 (2018).

47. Sussillo, D., Stavisky, S. D., Kao, J. C., Ryu, S. I. & Shenoy, K. V. Making brain–machine interfaces robust to future neural variability. Nat Commun 7, (2016).

48. Degenhart, A. D. et al. Stabilization of a brain–computer interface via the alignment of low-dimensional spaces of neural activity. Nature Biomedical Engineering 1–14 (2020) doi:10.1038/s41551-020-0542-9.

49. Radford, A. et al. Language models are unsupervised multitask learners. OpenAI Technical Report (2018).

50. Povey, D. et al. The Kaldi Speech Recognition Toolkit. IEEE 2011 Workshop on Automatic Speech Recognition and Understanding (2011).

51. Mohri, M., Pereira, F. & Riley, M. Speech Recognition with Weighted Finite-State Transducers. in Springer Handbook of Speech Processing (eds. Benesty, J., Sondhi, M. M. & Huang, Y. A.) 559–584 (Springer, 2008). doi:10.1007/978-3-540-49127-9_28.

