## Supplemental Figures and Tables for "High-performance brain-to-text communication via imagined handwriting"

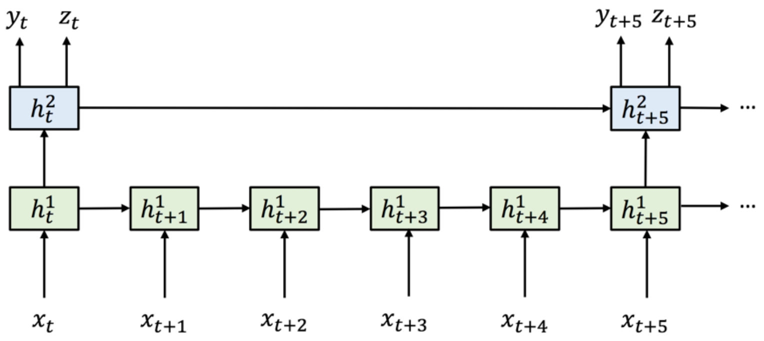


$x_t$ : binned, smoothed & z-scored firing rate vector (20 ms bins)

$h_t$ : hidden unit vector (512 units)

$y_t$: character probability vector

$z_t$: character start probability scalar

**GRU Update Equations**

$$r_t = \sigma(W_r x_t + R_h h_{t-1} + b_{Wr} + b_{Rr})$$
$$u_t = \sigma(W_u x_t + R_u h_{t-1} + b_{Wu} + b_{Ru})$$
$$c_t = \sigma_h(W_h x_t + r_t * (R_h h_{t-1} + b_{Rh}) + b_{Wh})$$
$$h_t = (1 - u_t) * c_t + u_t * h_{t-1}$$

**Output Equations**

$$y_t = \mathrm{softmax}(W_y h_t + b_y)$$
$$z_t = \sigma(W_z h_t + b_z)$$

**Supplemental Figure 1. Diagram and equations of our RNN architecture.** We used a two-layer gated recurrent unit (GRU) recurrent neural network architecture. The top layer runs at a slower frequency than the bottom layer, which we found improved the speed of training (by making it easier to hold information in memory for long time periods).

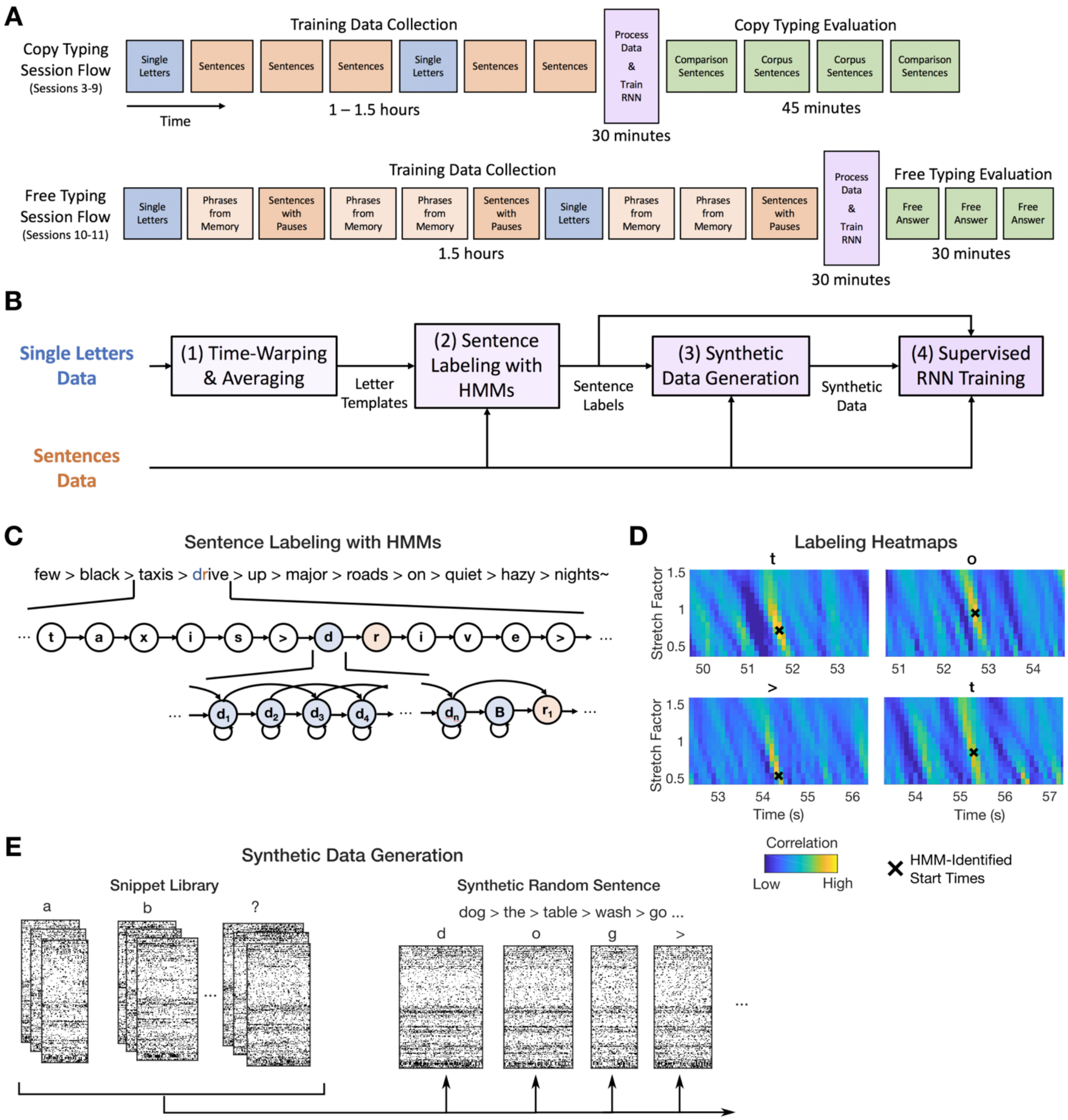


**Supplemental Figure 2. Overview of RNN training methods.** (A) Diagram of the session flow for copy typing and free typing sessions (each rectangle corresponds to one block of data). First, single letter and sentences training data is collected (blue and red blocks). Next, the RNN is trained using the newly collected data plus all previous days' data (purple block). Finally, the RNN is held fixed and evaluated (green blocks). (B) Diagram of the data processing and RNN training process (purple block in A). First, the single letter data is time-warped and averaged to create spatiotemporal templates of neural activity for each character. These templates are used to initialize the hidden Markov models (HMMs) for sentence labeling. After labeling, the observed data is cut apart and rearranged into new sequences of characters to make synthetic sentences. Finally, the synthetic sentences are combined with the real sentences to train the RNN. (C) Diagram of a forced-alignment HMM used to label the sentence “few black taxis drive up major roads on quiet hazy nights”. The HMM states correspond to the sequence of characters in the sentence. (D) The label quality can be verified with cross-correlation heatmaps made by correlating the single character neural templates with the real data. The HMM-identified character start times form clear hotspots on the heatmaps. (E) To generate new synthetic sentences, the neural data corresponding to each labeled character in the real data is cut out of the data stream and put into a snippet library. These snippets are then pulled from the library at random, stretched/compressed in time (to add more artificial timing variability), and combined into new sentences.

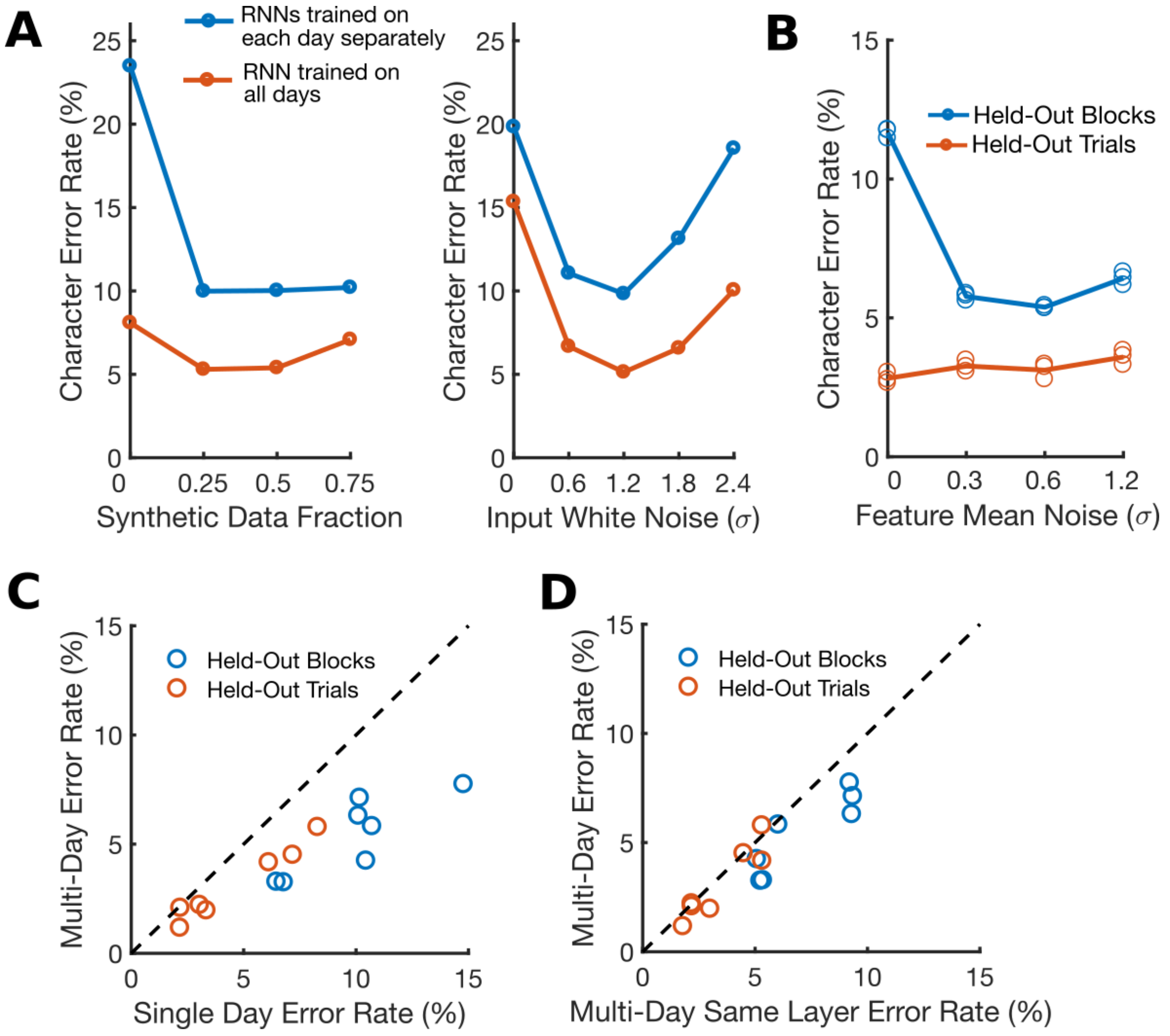


**Supplemental Figure 3. The effect of key RNN parameters on performance.** (A) Training with synthetic data (left) and artificial white noise added to the inputs (right) were both essential for high performance. Data are shown from a grid search over both parameters, and lines show performance at the best value for the other parameter. Results indicate that both parameters are needed for high performance, even when the other is at the best value. Using synthetic data is more important when the dataset size is highly limited, as is the case when training on only a single day of data (blue line). Note that the input white noise is in units of feature standard deviations (since the inputs given to the RNN were z-scored). (B) Artificial noise added to the feature means greatly improves the RNN's ability to generalize to new blocks of data that occur later in the session, but does not help the RNN to generalize to new trials within blocks of data that it was already trained on. This is because feature means change slowly over time. (C) Training an RNN with all of the datasets combined improves performance relative to training on each day separately. Each circle shows the performance on one of seven days. (D) Using separate input layers for each day is better than using a single layer across all days.

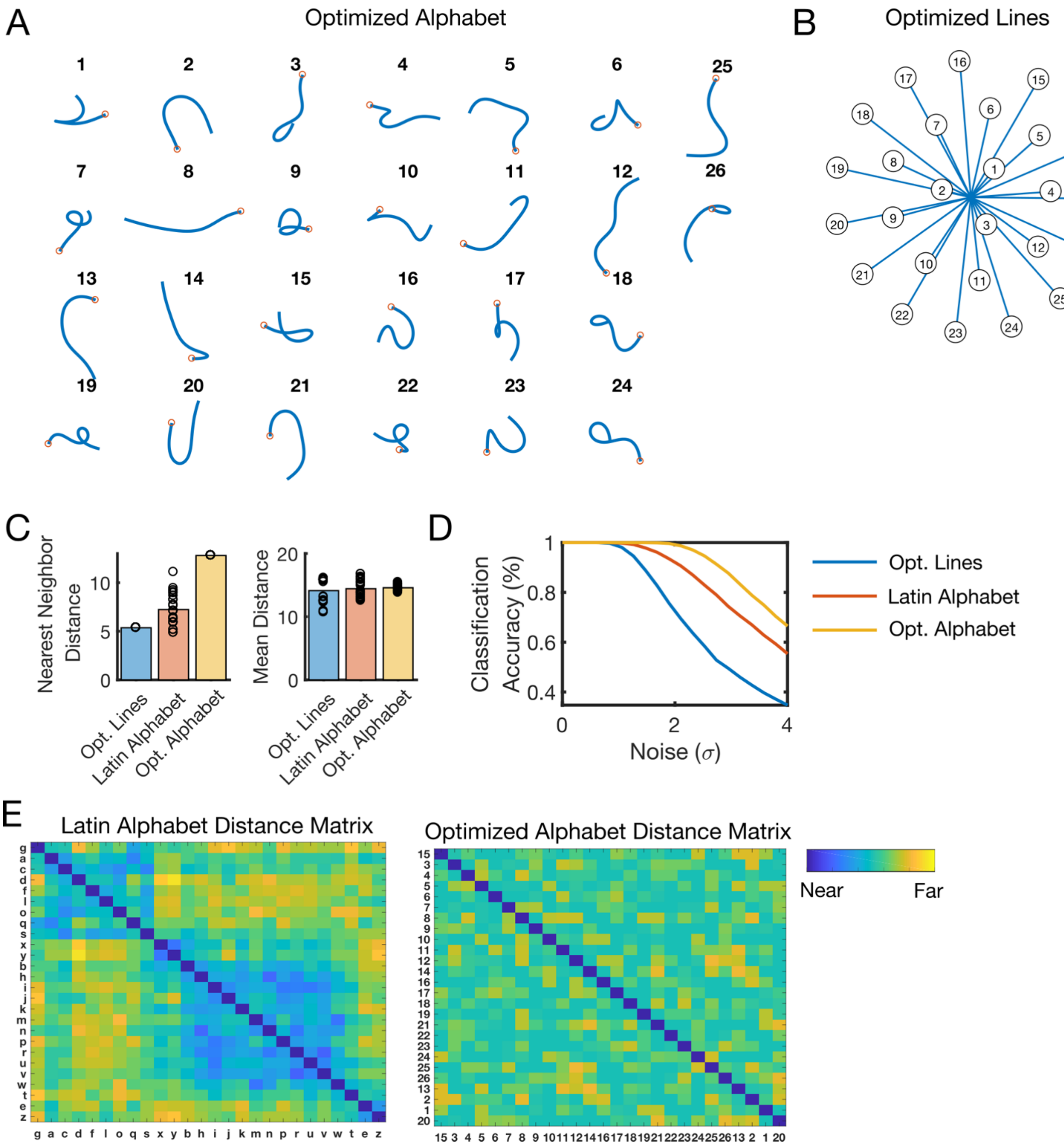


**Supplemental Figure 4. We optimized an artificial alphabet to maximize neural decodability.** (A) Using the principle of maximizing the nearest neighbor distance, we optimized for a set of pen trajectories that are theoretically easier to classify than the Latin alphabet (using standard assumptions of linear neural tuning to pen tip velocity). (B) For comparison, we also optimized for a set of 26 straight lines that maximize the nearest neighbor distance. (C) Pairwise Euclidean distances between pen tip trajectories were computed for each set, revealing a larger nearest neighbor distance (but not mean distance) for the optimized alphabet as compared to the Latin alphabet. Each circle represents a single movement and bar heights show the mean. (D)

Simulated classification accuracy as a function of the amount of artificial noise added. Results confirm that the optimized alphabet is indeed easier to classify than the Latin alphabet, and that the Latin alphabet is much easier to classify than straight lines, even when the lines have been optimized. (E) Distance matrices for the Latin alphabet and optimized alphabets show the pairwise Euclidean distances between the pen trajectories. The distance matrices were sorted into 7 clusters using single-linkage hierarchical clustering. The distance matrix for the optimized alphabet has no apparent structure; in contrast, the Latin alphabet has two large clusters of similar letters (letters that begin with a counter-clockwise curl, and letters that begin with a down stroke).

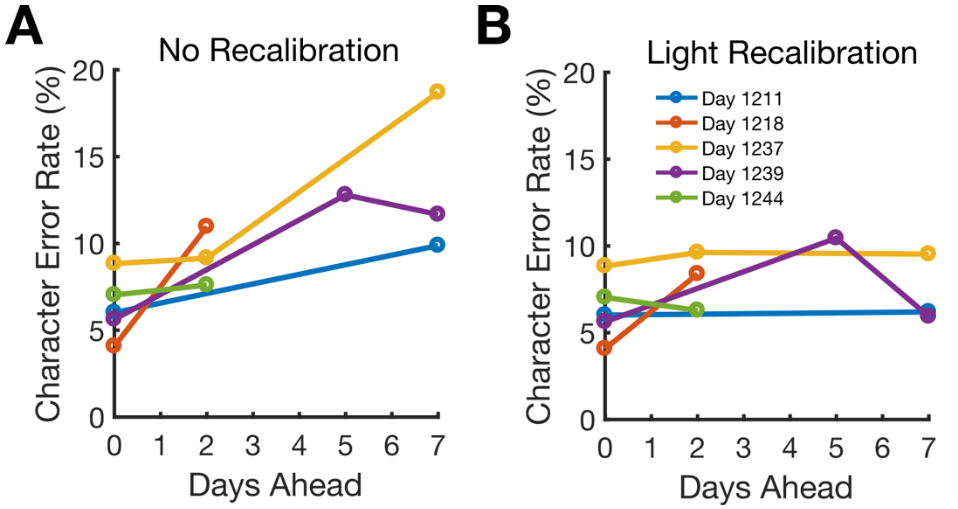


**Supplemental Figure 5. Cross-Day Transfer of RNNs.** (A) Performance of RNN decoders applied to future days without recalibrating. The decoders have some ability to transfer to new days (the decoder performance declines, but the error rates are still below 20% a week into the future). (B) If some "light" recalibration is performed (using 2 repetitions of each single character and 8 sentences, as opposed to 10 repetitions and 50 sentences), the performance decline can largely be prevented, suggesting that the full calibration routine we employed may not be necessary for achieving high performance. When doing light recalibration, we allowed only the input layer of the RNN to change, which we found protected against overfitting to the limited data. To further protect against overfitting, we only ran 200 minibatches of training (as opposed to our typical 1000).

| **Session #** | **Date (Trial Day)** | **Description** | **Data** |
| --- | --- | --- | --- |
| **1** | 2019.05.08 (994) | Sentence-writing and character-writing pilot day (no decoding) | Sentence writing collected as training data (102 sentences, no decoding); single character writing (27 repetitions per character) |
| **2** | 2019.06.03 (1020) | Attempted handwriting of straight lines | Instructed delay straight-line writing (24 repetitions each for 16 straight-line conditions) |
| **3** | 2019.11.25 (1195) | Real-time decoding pilot day | Sentence writing for training data (50 sentences, no decoding); sentence writing for performance evaluation (34 sentences, with real-time decoding); single character writing (10 repetitions per character). |
| **4** | 2019.12.09 (1209) | Real-time decoding pilot day | Sentence writing for training data (50 sentences, no decoding); sentence writing for performance evaluation (34 sentences, with real-time decoding); single character writing (10 repetitions per character). |
| **5** | 2019.12.11 (1211) | Copy-typing evaluation | Sentence writing for training data (50 sentences, no decoding); sentence writing for performance evaluation (34 sentences, with real-time decoding); single character writing (10 repetitions per character). |
| **6** | 2019.12.18 (1218) | Copy-typing evaluation | Sentence writing for training data (50 sentences, no decoding); sentence writing for performance evaluation (34 sentences, with real-time decoding); single character writing (10 repetitions per character). |
| **7** | 2019.12.20 (1220) | Copy-typing evaluation | Sentence writing for training data (50 sentences, no decoding); sentence writing for performance evaluation (34 sentences, with real-time decoding); single character writing (10 repetitions per character). |
| **8** | 2020.01.06 (1237) | Copy-typing evaluation | Sentence writing for training data (50 sentences, no decoding); sentence writing for performance evaluation (34 sentences, with real-time decoding); single character writing (10 repetitions per character). |
| **9** | 2020.01.08 (1239) | Copy-typing evaluation | Sentence writing for training data (50 sentences, no decoding); sentence writing for performance evaluation (34 sentences, with real-time decoding); single character writing (10 repetitions per character). |
| **10** | 2020.01.13 (1244) | Free-answer evaluation | Sentence writing with artificial pauses (30 sentences, no decoding); phrase writing from memory (200 phrases, no decoding); sentence writing for performance evaluation (25 free-answer questions); single character writing (10 repetitions per character). |
| **11** | 2020.01.15 (1246) | Free-answer evaluation | Sentence writing with artificial pauses (30 sentences, no decoding); phrase writing from memory (200 phrases, no decoding); sentence writing for performance evaluation (25 free-answer questions); single character writing (10 repetitions per character). |

**Table S1.** List of all data collection sessions included in this study.

| Prompt | Real-Time Output | Real-Time Output (Annotated) | Retrospective Language Model (Annotated) |
|---|---|---|---|
| **infected adults develop a cough and their skin and ears turn blue.** | infected adults dercep a cough and thhir skin and ears tunn blue | infected adults dercep a cough and thhir skin and ears tunn blue. | infected adults develop a cough and their skin and ears turn blue. |
| **i interrupted, unable to keep silent.** | i interrupted, unabee to keep silent. | i interrupted, unabee to keep silent. | i interrupted, unable to keep silent. |
| **i dumped the tools in the hut.** | i dumped the tools in the hut. | i dumped the tools in the hut. | i dumped the tools in the hut. |
| **within thirty seconds the army had landed.** | within thirty seconds the armmy had landed | within thirty seconds the armmy had landed. | within thirty seconds the army had landed. |
| **that's when i threw up on the carpet.** | that's when i threw up on the carpet. | that's when i threw up on the carpet. | that's when i threw up on the carpet. |
| **he didn't want to rub salt into her wounds.** | he didn't want to rub salt into her wounds. | he didn't want to rub salt into her wounds. | he didn't want to rub salt into her wounds. |
| **shouting and swearing, i yelled for an epidural.** | shouting and sweerin, I yeled for an epidural. | shouting and sweerin~~g~~, i yel~~l~~ed for ■ an epidural. | shouting and swearing, i yelled for a an epidermal. |
| **you wish to purchase something?** | you wish to purchase something? | you wish to purchase somathing? | you wish to purchase something? |
| **lowell felt like a soldier on a battlefield, stripped of ammunition.** | lowel felt like a soldier on a battlefield, stripped ef ammunition. | lowel~~l~~ felt like a soldier on a battlefield, stripped ef ammunition. | lowell felt like a soldier on a battlefield, stripped of ammunition. |
| **there are only one or two minor casualties.** | thee ane only on or two minor cafualties. | the~~r~~e ane only one or two minor cafualties. | there are only one or two minor casualties. |

**Table S2.** Example decoded sentences for one block of copy typing. In the rightmost columns, errors are highlighted in red ( ■ indicates an extra space, omitted letters were indicated with a strikethrough). Note that our language model substitutes "epidermal" for "epidural", since "epidural" is out of vocabulary. The mean characters per minute for this block was 86.47 and the character error rates were 4.18% (real-time output) and 1.22% (language model).

| Prompt | Real-Time Output | Real-Time Output (Annotated) | Retrospective Language Model (Annotated) |
|---|---|---|---|
| **what made you first get interested in machining?** | iit was the famil business | iit was the famil~~y~~ business | it was the family business |
| **what is the hardest part of machining?** | forming the tooling | forming the tooling | forming the tooling |
| **how much spice do you like in your food?** | lots and lots | lots and lots | lots and lots |
| **what type of music do you enjoy most?** | ii like loud rock musica | ii like loud rock musica | i like loud rock music |
| **what are some of your favorite games to play?** | i like to play cards | i like to play cards | i like to play cards |
| **what has taken you the longest to get good or decent at?** | i worked for years to perfecet my photography | i worked for years to perfecet my photography | i worked for years to perfect my photography |
| **what food do you love that a lot of people might find a little odd?** | very few poope can eat smeked sailfish | very few poop~~l~~e can eat smeked sailfish | very few people can eat smoked salt fish |
| **if you could start a charity, what would it be for?** | i woud proovide furding to smal families | i woud proovide furding to smal~~l~~ families | i would provide funding to small families |
| **what advice would you give to your younger self?** | be patientt it wil got better | be patientt it wil~~l~~ got better | be patient, it will get better |

**Table S3.** Example decoded sentences for one block of free typing. In the rightmost columns, errors are highlighted in red (omitted letters were indicated with a strikethrough). Note that our language model substitutes "salt fish" for "sailfish", since "sailfish" is out of vocabulary. The mean characters per minute for this block was 73.8 and the estimated character error rates were 6.82% (real-time output) and 1.14% (language model).

| | Held-Out Trials + Mean Subtraction | Held-Out Trials | Held-Out Blocks |
|---|---|---|---|
| **RNN** | 0.23 % | 0.23 % | 0.70 % |
| **HMM** | 2.96 % | 6.70 % | 80.08 % |

**Table S4.** Offline performance (character error rate) of the RNN decoder compared to a hidden Markov model (HMM) decoder, both with a language model applied. Results show that the RNN strongly outperforms the HMM, especially in situations where the feature means are likely to have changed. The HMM has no built-in mechanism for adapting to changes in the feature means, leading to very poor performance when generalizing to held-out blocks, and best performance when subtracting within-block feature means to account for any changes that may have occurred over time.

**Supplemental Video Legends**

**Supplemental Video 1. Copying sentences in real-time with the handwriting brain-computer interface.** In this video, our participant (T5) copies sentences displayed on a computer monitor with the handwriting-brain computer interface. When the red square on the monitor turns green, this cues T5 to begin copying the sentence.

**Supplemental Video 2. Freely answering questions in real-time with the handwriting brain-computer interface.** In this video, our participant (T5) answers questions that appear on a computer monitor using the handwriting brain-computer interface. T5 was instructed to take as much time as he wanted to formulate an answer, and then to write it as quickly as possible.

**Supplemental Video 3. Side-by-side comparison between the handwriting brain-computer interface and the prior state of the art for intracortical brain-computer interfaces.** In a prior study (Pandarinath et al., 2017) our participant achieved the highest typing speed ever reported with an intracortical brain-computer interface (39 correct characters per minute using a point-and-click typing system). Here, we show an example sentence typed by our participant using the point-and-click system (shown on the bottom) and our new handwriting brain-computer interface (shown on the top), which is more than twice as fast.

**Supplemental Video 4. Hand micromotion while using the handwriting brain-computer interface.** Our participant is paralyzed from the neck down (C4 ASIA C spinal cord injury) and only generates small micromotions of the hand when attempting to handwrite.
