## Supplemental Methods for "High-performance brain-to-text communication via imagined handwriting"

### Supplemental Materials

### RNN Training Procedure

The RNN training procedure consists of four stages, as diagrammed in SFig. 2B. The training procedure was designed to address two main challenges: (1) inferring when each character was written in the training data, so that supervised learning techniques could be used to train the RNN, and (2) augmenting the training data with synthetic sentences, to prevent the RNN from overfitting to limited data.

#### Data Preprocessing

The single character data was pre-processed by binning the recorded threshold crossings into 10 ms bins. The binned rates were then z-scored (by subtracting the mean of each electrode and dividing by its bin-by-bin standard deviation) and smoothed by convolving with a gaussian kernel (sd = 30 ms).

The sentence data was also binned (10 ms bins), smoothed by convolving with a gaussian kernel (sd = 40 ms), and z-scored. For the sentence data, z-scoring was performed by applying the means and standard deviations from all trials of the single character data. During real-time decoding, these same means and standard deviations were also used to z-score the data.

#### Stage 1: Single Character Time Warping & Averaging

For each session, at least two blocks of single character data were collected as part of the training data (10 repetitions of each character total). We then used time-warped PCA (https://github.com/ganguli-lab/twpca)(Poole et al., 2017; Williams et al., 2020) to find continuous, regularized time-warping functions to align all trials corresponding to a single character together. We used the following time-warping parameters: 5 components (temporal factors), 0.001 scale warping regularization (L1), and 1.0 scale time regularization (L2).

After time-warping, the neural activity was averaged across trials for each character, yielding an N x T mean neural activity matrix for each character. N is the number of microelectrodes (192) and T is the number of time steps. The time window to use for each character was chosen by visual inspection of the character shapes, using the data from session 1 (i.e., the shapes shown in Figure 1). After the character durations were chosen, they were held fixed for all subsequent sessions. The first 100 ms of reaction time after the go cue was excluded, yielding a time window for each character that spanned 100 ms after the go cue until the end time chosen by inspection. Finally, the neural activity was then downsampled to 50 ms time steps by averaging every 5 bins together, yielding single character “neural templates” that were used to build HMMs for data labeling (see below).

#### Stage 2: Sentence Labeling with Hidden Markov Models

Data labeling was accomplished in 6 steps as follows:

(1) Construct an HMM for each sentence, using the neural templates from the single character data.
(2) Use the Viterbi algorithm to infer when each character began and ended in each sentence.
(3) Refine the character start times and durations using a local grid search.
(4) Update the HMM emission probabilities (but not the state transition probabilities) using the inferred character start times and durations.
(5) Repeat Steps 2-3.
(6) Construct the final targets for supervised RNN training ($y_t$ and $z_t$), using the letter start times found in (5).

Step 1: HMM Construction: We used hidden Markov models (HMMs) to label our data, similar to how HMMs have been used in speech recognition to determine when phonemes begin and end in an utterance where the transcription is known (this is called "forced alignment") (Young et al., 2006). For each sentence of training data, we constructed an HMM whose states and state transitions defined an orderly march through the characters of that sentence (SFig. 2C). Each individual character was represented with a sequence of HMM states, whose multivariate Gaussian emission probabilities were determined by the single character neural templates. Specifically, for each character, the number of HMM states was equal to the number of 50 ms bins in the neural template for that character (plus an additional "blank" state), and the mean vector for each state's emission distribution was equal to the corresponding firing rate vector from the neural template. The covariance matrix was set equal to the identity matrix.

Let us denote the states of the HMM model as $s_{i,j}$, where $j$ denotes the character number in the sentence, and $i$ iterates through the states within each character. The state transitions for states $s_{1,j}$ to $s_{N,j}$ (and the optional blank state $B_j$) for each character were hand-set to the following reasonable values (where $N$ is the number of states in character $j$, and $M$ is the total number of characters in the sentence):

| States | Description | Transition Probabilities |
|---|---|---|
| $s_{x,j}$ for x<N-1 | All states before the second to last, for character *j* | $P(s_{x,j} \rightarrow s_{x,j}) = 0.2$<br>$P(s_{x,j} \rightarrow s_{x+1,j}) = 0.6$<br>$P(s_{x,j} \rightarrow s_{x+2,j}) = 0.2$ |
| $s_{N-1,j}$ | Second to last state for character *j* | $P(s_{N-1,j} \rightarrow s_{N-1,j}) = 0.2$<br>$P(s_{N-1,j} \rightarrow s_{N,j}) = 0.8$ |
| $s_{N,j}$ | Last state for character *j* | $P(s_{N,j} \rightarrow s_{N,j}) = 0.2$<br>$P(s_{N,j} \rightarrow B_j) = 0.1$<br>$P(s_{N,j} \rightarrow s_{1,j+1}) = 0.7$ |
| $B_j$ | Blank state for character *j* | $P(B_j \rightarrow B_j) = 0.5$<br>$P(B_j \rightarrow s_{1,j+1}) = 0.5$ |
| $s_{N,M}$ | Last character state in the sentence | $P(s_{N,M} \rightarrow s_{N,M}) = 0.7$<br>$P(s_{N,M} \rightarrow B_M) = 0.3$ |

| | | |
|---|---|---|
| $B_M$ | Last blank state in the sentence | $P(B_M \rightarrow B_M) = 1.0$ |

Note that $B_j$ is a "blank" state that can be entered into at the end of the $j$th character (or skipped) and can model pauses in between characters. The emission vector for all blank states was set to the average neural activity vector across all character templates.

Step 2: Viterbi Algorithm We used the Viterbi algorithm (Rabiner, 1989) to find the most likely sequence of HMM states given the observed neural activity, with the constraint that the last state of the sequence was the last character's final state ($s_{N,M}$) or the last blank state ($B_M$) (this enforces that the sentence is completely finished). We added an additional constraint that helped prevent pathological solutions: each state had to occur within a certain window of time centered on its character's location in the sentence. Let us denote the duration of the entire sentence as $T$. Each character $j$'s time window was centered on the time $(j/M)T$ and extended $0.3T$ in either direction. For example, if the character 'm' occurred in the middle of the sentence, then the states for this 'm' had to occur between times $0.2T$ and $0.8T$. Similarly, if the character 't' occurred in the beginning of the sentence, it had to occur between times 0 and $0.3T$. This constraint was implemented by setting a states' observation probability to negative infinity if it lied outside of this window.

Step 3: Local Refinement of Character Start Times The sequence of states found by the Viterbi algorithm define the start time and duration of each character. The quality of fit can be roughly assessed with correlation heatmaps that show the correlation (Pearson's $r$) between the neural template for a character and the observed neural activity, as a function of character start time and character stretch factor (SFig. 2D). The identified start time and duration should lie on a hotspot. For these heatmaps, the correlation coefficient was computed for each microelectrode channel separately; the resulting 192 correlation coefficients were then averaged together to produce a final value.

We implemented a refinement step after the Viterbi search which maximized the correlation of each character with the observed activity via a grid search of adjusted start times and template stretch factors. The grid search varied the possible start times from 0.5 seconds before to 0.5 seconds after the HMM-identified time (in steps of 0.05 seconds). The stretch factor varied from 0.4 to 1.5 in steps of 0.0786. Values that caused the adjusted character template to intersect adjacent character templates were not considered. This refinement procedure effectively placed each character on a nearby maximum in the heatmaps shown in SFig. 2D.

Step 4: Updating HMM Emissions We updated the HMM emission probabilities based on the newly labeled sentence data. The emission probabilities were updated in the following way. First, for each character class, all example snippets of that character were gathered together based on the character start times and durations found above. Then, each example was time-normalized by resampling to N time steps (using linear interpolation, where N was the original number of time steps in the template). Then, the time-normalized examples were averaged together to compute a new neural template for that character. The emission probabilities were not updated for characters with less than 18 examples (e.g., rare characters such as "q" or "x").

In principle, the state transition probabilities of the HMM could also be updated (e.g. by using the Baum-Welch algorithm). However, we did not explore that here, as we found that updating the emission probabilities alone seemed sufficient to yield high quality labels.

Step 5: Repeat with New HMM Emissions With the updated emissions, we performed one additional iteration of HMM labeling and subsequent refinement (further iterations did not seem to improve label quality).

Step 6: Construct RNN Targets Finally, target variables for supervised RNN training were generated using the letter start times found above. Two target time series were created: a series of one-hot character vectors ($y_t$), where each vector is a one-hot representation of the most recently started character, and a scalar time series ($z_t$) that indicates whether *any* new character has recently been started. The $z_t$ signal allows repeated characters to be distinguished (these would otherwise appear identical to a longer, single character as seen through $y_t$).

Intuitively, $y_t$ is a "sample and hold" signal that stores whatever the most recently started character was indefinitely. For example, even if T5 pauses for several seconds after writing the character "a", $y_t$ will still continue to reflect "a" indefinitely until a new character is started. The $z_t$ signal is a complementary binary signal that goes high for a brief time whenever *any* new character begins. $z_t$ can be thresholded to detect the presence of new letters and type them on the screen, which we did online. More formally, $y_t$ and $z_t$ were defined as follows:

$$y_{t,i} = \begin{cases} 0, & \text{the most recently started character was not i} \\ 1, & \text{the most recently started character was i} \end{cases}$$

$$z_t = \begin{cases} 0, & \text{the most recent character was started} > 200 \text{ ms ago} \\ 1, & \text{the most recent character was started} \leq 200 \text{ ms ago} \end{cases}$$

One potential advantage of this nontraditional representation is that only the character start times are required; thus, any uncertainty about when each character ends shouldn't degrade performance (i.e., uncertainty about the length of time spent transitioning between letters, either with long pauses or short bouts of pen repositioning). Additionally, by not including multiple sub-states per character (which could be an alternative way to distinguish repeated letters), this method gives the RNN freedom to decide how to break apart each character into sub-states.

#### Stage 3: Synthetic Data Generation

Once the character start times were inferred for each sentence, we generated new synthetic sentences by rearranging the characters into different sequences (SFig. 2E). This data augmentation step was essential for high performance (SFig. 3A), as it helped to prevent the

RNN from overfitting to a limited training dataset. Here, we describe the synthesis process in detail.

Making the Snippet Library: First, we created a snippet library of neural activity snippets for each character. Entries were taken by extracting time windows of activity from the sentences data, beginning at the letter start time identified by the HMM procedure and ending at the start time of the next letter. In this way, any pauses and transition-related activity are included at the end of the snippets.

Generating Random Sentences: Next, we used the snippet library to generate synthetic sentences (24 seconds long, which was the length of data used in each minibatch during RNN training). First, the character sequence for each sentence was chosen by selecting words at random from a list of 10,000 common words (the 10,000 most frequent words appearing in the Google Web 1T 5-gram database) (Brants and Franz, 2006). Words were selected one at a time, with no dependence on the prior word, according to the following simple heuristic:

- 64% chance: a word was chosen uniformly at random from the entire list
- 20% chance: one of the twenty most frequent words was chosen uniformly at random
- 16% chance: a word with rare letters ("q", "x", "j", or "z") was chosen uniformly at random from the set of all such words (this helped prevent the RNN from neglecting rare characters)

To make sure punctuation characters were represented, apostrophes were randomly added in between the last and second-to-last letter of the word (3% chance) and commas were randomly added at the end of the word (7% chance). All words either ended in a period (5% chance), question mark (5% chance), or space (90% chance).

We used the above heuristic to generate sentences instead of using real sentences in order to discourage the RNN from "baking-in" a model of the English language that extends beyond single words.

Synthesizing the Neural Activity: Once the synthetic character sequence was determined, the corresponding neural activity was synthesized one character at a time. For each character, a snippet was chosen from the library at random in a way that attempted to respect pen transition movements between letters. For example, when transitioning from 'e' to 't', the pen must traverse upwards before beginning the downstroke for 't'. However, when transitioning from 'd' to 't', no such pen re-positioning is needed (when written in the way shown in Figure 1). To do this, we discretized the starting heights for each character to the following values: 0, 0.25, 0.5, 1. The assignment of each letter to each category is depicted in the table below.

| Start Height | 0 | 0.25 | 0.5 | 1.0 |
|---|---|---|---|---|
| Character | comma | a, o, e, g, q | c, d, m, j, i, n, p, r, s, u, v, w, x, y, z, space (>), period (~) | b, t, f, h, k, l, apostrophe, question mark |

When choosing a snippet from the library, we selected at random from all snippets whose next character in the training data began at the same height as the next character in the synthetic sentence. When this wasn't possible, we selected uniformly at random from all snippets.

After a snippet was chosen, we randomly time-warped the snippet by resampling it to a different length of time (chosen uniformly from 0.7 to 1.3 times its original length). This helps the RNN to be more robust to changes in letter timing. Finally, we also sometimes added a long pause at the end of the snippet at random, to train the RNN to be robust to unpredictable pauses made by the user. The probability of adding a pause was 3%; if a pause was added, its duration was drawn from an exponential distribution (mean of 1 second). The synthetic neural activity during the pause was white noise with a standard deviation of 1.

#### Stage 4: Supervised RNN Training

The RNN architecture is described in the Methods and illustrated in SFig. 1. Here, we describe in detail how the RNN weights were optimized using supervised learning.

Implementation: Optimization of the RNN weights was implemented with TensorFlow v1.15 (Abadi et al., 2016). We used a desktop machine with 4 NVIDIA GeForce GTX 1080 Ti GPUs and a 32-core AMD RYZEN Threadripper 2990WX CPU to train the RNNs. To train a single RNN, only a single GPU was used, allowing parallel training of up to 4 RNNs. A single minibatch took ~0.25 seconds to complete, resulting in training times ranging from 4 minutes (1k minibatches, when updating the RNN to a new day of data) to 3.5 hours (50k minibatches, when training from scratch). Note that the number of sentences used for training was small (572 by the last day of copy typing), so only a few minibatches were required to cycle through all sentences.

Mini-batches & gradient descent: Gradients were computed on mini-batches of 64 data snippets using backpropagation through time (Goodfellow et al., 2016). We used the gradient descent method "Adam" (Kingma and Ba, 2017) (beta1 = 0.9, beta2 = 0.999, epsilon = 0.01). The learning rate was decreased linearly from 0.01 to 0 over a pre-specified number of minibatches (1k when updating a pre-trained RNN with a new day of data, 50k when training from scratch). To prevent gradient explosion, gradient magnitudes were clipped at 10.

Each data snippet used in a mini-batch was 24 seconds long and was selected at random from the training sentences by uniformly drawing a random start time from 0 to $\tau$-24, where $\tau$ is the duration of the sentence. Each minibatch mixed together synthetic sentences and real sentences, in a proportion that was tuned to optimize performance (beginning at 75% synthetic and ending at ~40%). Each mini-batch selected data snippets from a single day only, which was chosen at random amongst all available days of training data. We weighted the most recent day more highly.

Cost Function: We used the following cost function for a single snippet of data, which expresses the sum of an L2 weight regularization, a cross-entropy loss, and a squared error loss:

$$\lambda \sum_i \|W_i\|_F^2 - \frac{1}{T}\sum_{t=50}^{T}\sum_{c=1}^{C} y_{t,c}\log \hat{y}_{t+d,c} + \frac{1}{T}\sum_{t=50}^{T}(z_t - \hat{z}_{t+d})^2$$

Here, λ scales the L2 regularization of the RNN weight matrices $W_i$ (penalizing large weights), $T$ is the number of time steps in the data snippet (1200, 20 ms time steps), $C$ is the number of characters (31), $y_{t,c}$ is a one-hot representation of the most recently started character at time step $t$, $\hat{y}_{t+d,c}$ is the RNN's prediction of $y_{t,c}$ ($d$ time steps in the future, d=50), $z_t$ is a scalar representation of whether a character was started within the last 200 ms, and $\hat{z}_{t+d}$ is the RNN's prediction of $z_t$.

The one second delay ($d$) between the RNN output ($\hat{y}_{t+d,c}$ , $\hat{z}_{t+d}$ ) and the target signals ($y_{t,c}$, $z_t$) was added to give the RNN enough time to observe all of the neural activity corresponding to a character before deciding on its identity. Note also that there is a one second "burn-in" time before the error is counted, to ensure that the RNN is not penalized for incorrectly identifying characters at the very beginning of the snippet that may begin somewhere in the middle of the character (since snippets start at random times).

Noise Perturbations: We added two types of artificial noise to the neural features in order to regularize the RNN. First, we added white noise directly to the input feature vectors, which greatly improved performance (SFig. 3A, middle panel). Adding white noise to the inputs asks the RNN to map clouds of similar inputs to the same output, improving generalization. The standard deviation of the white noise was an important hyperparameter that we tuned to optimize performance (see parameter values below).

We also added artificial changes to the means of the neural features, to make the RNN robust to non-stationarities in the neural data (which has been an important problem for intracortical BCIs (Jarosiewicz et al., 2015; Sussillo et al., 2016)). These artificial mean changes greatly improved the RNN's ability to generalize to held-out blocks of data occurring later in a session (SFig. 3B). We added two types of perturbations to the neural features to simulate non-stationarities: constant offset noise and random walk noise.

The above-mentioned types of noise (white noise, constant offset noise, and random walk noise) were all combined together to transform the input vector in the following way:

$$\tilde{x}_t = x_t + \varepsilon_t + \varphi + \sum_{i=1}^{t} \upsilon_i$$

Here, $x_t$ are the original neural features, $\varepsilon_t$ is a white noise vector unique to each time step, $\varphi$ is a constant offset vector, and $\upsilon_t$ are white noise vectors that are cumulatively summed to simulate a random walk.

Combining Data Across Days: Combining multiple days of data together greatly improved performance relative to using just a single day (SFig. 3C). To combine data across days, we optimized day-specific affine transforms of the input that could account for changes in the neural features across days (as observed previously in e.g. (Jarosiewicz et al., 2015; Downey et al., 2018; Degenhart et al., 2020)):

$$\acute{x}_t = A_i \tilde{x}_t + b_i$$

Here, $\tilde{x}_t$ is a vector of neural features (after artificial noise has been added, see above), $A_i$ is a 192 x 192 matrix, and $b_i$ is a 192 x 1 vector. The transformed features were then fed as input to the RNN. The $A_i$ and $b_i$ parameters are optimized simultaneously along with all other RNN parameters. Using separate input layers for each day improved performance relative to using a single shared input layer (SFig. 3D).

Hyperparameters: We summarize the hyperparameters used for RNN training in the table below. Most parameters were hand-tuned; in later days, we performed small hyperparameter optimization routines where we trained 100 RNNs with parameters drawn at random (from pre-specified lists of reasonable values). Parameters for the next session were then taken from the top performing RNN. Thus, sometimes different hyperparameter values were used on different sessions; in particular, the amount of regularization decreased as more data were accumulated (the fraction of synthetic sentences and the white noise became smaller). Since parameters varied, we summarize their typical ranges and values (rather than exhaustively list each combination for each session).

| Parameter | Description | Typical Value(s) |
|---|---|---|
| λ | L2 Weight Regularization | 1e-5 |
| H | Hidden state size | 512 |
| σ | Standard deviation of white noise | 1.0 – 1.6 (SFig. 3A) |
| γ | Standard deviation of constant offset noise | 0.6 (SFig. 3B) |
| ζ | Standard deviation of random walk noise | 0.02 |
| χ | Fraction of synthetic trials used in each mini-batch | 0.375 – 0.75 (SFig. 3A) |
| ω | Probability of a min-batch drawing sentences from the most recent day | 0.25 – 0.5 |
| α | Learning rate | 0.01 |
| M | Number of min-batches to train | 1k (updating for new day), 50k (training from scratch across all days) |
| N | Min-batch size | 64 |
| T | Duration of data snippets in each mini-batch | 24 seconds |

### Language Model

#### WebText Preprocessing

Our bigram language model was created using 250k samples of text from WebText (https://github.com/openai/gpt-2-output-dataset), provided by OpenAI (San Francisco, CA). We pre-processed the WebText samples to convert them to lowercase, remove symbols that were not in our character set, and split into sentences (yielding a total of 5.1M sentences). We applied the following step-by-step recipe to pre-process the sentences:

(1) Replace newlines and hyphens with spaces
(2) Convert all letters into lower-case
(3) Delete all characters not in the character set (which consisted only of the letters a-z, periods, spaces, commas, question marks and apostrophes)
(4) Replace repeated spaces with single spaces
(5) Remove spaces in front of periods and commas
(6) Replace repeated periods with single periods
(7) Strip surrounding whitespace from the sample
(8) Split the sample into sentences by splitting at periods and question marks

#### Constructing the Bigram Language Model

We used publicly available scripts [(Puigcerver, 2017); https://github.com/jpuigcerver/Laia/tree/master/egs/iam] as a starting point for constructing our bigram language model. These scripts tokenize the list of sentences created above into discrete words, and then call Kaldi (Povey et al., 2011) and SRILM (Stolcke et al., 2011) programs to count the frequency with which these words (and pairs of words) appear in the sentences. Word frequencies are then encoded into a weighted finite state transducer (Mohri et al., 2008) that can be used to infer the most likely sequence of characters given the word frequency counts and the character probabilities from the neural decoder.

To ensure that our language model was sufficiently general to allow the expression of a wide variety of sentences, we used a large vocabulary (consisting of the 50,000 most common words in the WebText samples). A space character was included at the beginning of each word to enforce that words were always separated by spaces (except for words containing punctuation, such as contractions).

In big picture, the language model is essentially a large hidden Markov model where each state corresponds to a character, and where the state transition probabilities encode the statistics of which words are likely to follow other words. Inference is done with the language model by using an approximate Viterbi search (beam search) to find likely sequences of characters. The beam search combines information from the language priors (state transition probabilities) and information from the neural decoder about which characters are likely occurring at each moment in time (observations).

The language model was represented as the composition of weighted finite state transducers (Mohri et al., 2008) that encode information about different parts of the model:

$$H \circ C \circ L \circ G$$

Here, G is the grammar that encodes legal sequences of words and their probabilities (based on the unigram and bigram probabilities), L is the lexicon that encodes what characters are contained in each legal word, and H and C encode information about the character sub-states used in decoding (see Kaldi documentation for more detail). Each character had two sub-states $s_1$ and $s_2$. $s_1$ emits a CTC blank [(Puigcerver, 2017) used a neural network trained with the CTC loss function, see below] and $s_2$ emits the corresponding character. We used the following transition probabilities: P($s_1 \rightarrow s_1$)=0.6, P($s_1 \rightarrow s_2$)=0.2, P($s_1 \rightarrow s_{next}$)=0.2, P($s_2 \rightarrow s_2$)=0.6, P($s_2 \rightarrow s_{next}$)=0.4 , where $s_{next}$ is the first sub-state of the next character.

#### Inference with the Bigram Language Model

The scripts we used from Puigcerver and colleagues (Puigcerver, 2017) configured the language model to work with outputs generated by a neural network trained with the connectionist temporal classification (CTC) loss function (Graves et al., 2006). We therefore transformed our neurally decoded probabilities to make them look more like CTC outputs before using the language model. To generate the CTC blank probability $blank_t$, we transformed $z_t$ (the character start signal):

$$blank_t = 1 - \sigma(4 + 4\sigma^{-1}(z_{t+20}))$$

This hand-tuned function inverts $z_t$ and makes it sharper, so that it stays mostly at 1 and dips to 0 only briefly whenever a new character is written. It also shifts the signal forward by 20 time steps (400 ms), so that it dips to 0 at times when the character probabilities $y_t$ have already finished transitioning from the previous character to the next. Finally, we also modified $y_t$ so that all entries of $y_t$ plus the blank signal $blank_t$ sum to 1:

$$y_t' = y_t(1 - blank_t)$$

To perform inference with the language model and the above probabilities, we used Kaldi (and custom decoding functions by Puigcerver https://github.com/jpuigcerver/kaldi-decoders) to perform a beam search to generate lattices of candidate word sequences with high likelihood (beam=65, max active=5000, acoustic score=1.0, lattice beam=10). On average, the decoding process completed 3.74 times faster than real-time (averaged over the last two sessions of copy typing evaluation, where T5 wrote the fastest).

#### Rescoring with GPT-2

We used OpenAI's neural network language model "GPT-2" to rescore the candidate sentences inferred by the bigram language model [(Radford et al., 2018), 1558M parameter version, https://github.com/openai/gpt-2]. Rescoring using a neural network model is motivated by the fact that neural networks are powerful language models that can model long-range semantic dependencies, but may be too slow to use to efficiently search through many different possibilities. Thus, a simpler N-gram model can propose a list of plausible candidate sentences which a neural network model rescores [e.g., (Xiong et al., 2017)].

When run on a sequence on characters, GPT-2 returns the conditional probability of each character given the previous characters. These probabilities can be used to compute the log probability of any candidate sentence in the following way:

$$\log P(c_1, c_2, \dots, c_N) = \log P(c_1) + \log P(c_2 \mid c_1) + \dots + \log P(c_N \mid c_{N-1}, \dots, c_1)$$

Here, $P(c_1, c_2, \dots, c_N)$ is the probability of observing the character sequence $c_1, c_2, \dots, c_N$ and $P(c_N \mid c_{N-1}, \dots, c_1)$ is the conditional probability of observing character $c_N$ given previous characters $c_{N-1}, \dots, c_1$.

When generating candidate sentences for rescoring, an "acoustic score" and a "language model score" was returned for each candidate sentence. The acoustic score contains the cumulative log probability of observing the character sequence given the sentence, and the language model score contains the log probability of observing the sentence given the language model. When rescoring, we replaced the language model score with the probability returned by GPT-2, scaled the acoustic score by 0.5, and summed them together to generate a final score. The minimum score across all candidate sentences was then chosen.

#### Performance without rescoring

We compared the performance of the language model with rescoring to the bigram model alone. When decoding with the bigram model alone, we also used an acoustic score of 0.5. The character error rate with the bigram model alone was 1.48% [1.11, 1.85] (95% CI), and the word error rate was 4.72% [3.67, 5.87].

#### Bidirectional RNN

Bidirectional RNNs are used commonly whenever the computation at hand is not required to be causal (e.g. for non-real-time speech recognition, or for machine translation). To implement bidirectionality, we made each of the two layers (depicted in SFig. 1) bidirectional by adding an identical GRU component that runs in the opposite direction (i.e., begins with the last neural feature vector $x_T$ and ends with the first neural feature vector $x_1$). The first RNN layer thus has a total of 1024 hidden units (512 units in the forward direction, 512 units in the backwards direction); these 1024 hidden units were concatenated together in a vector and fed as input to both the forward and backward components of the second layer. Likewise, the hidden state of both the forward and backward components in the second layer were concatenated together at each time step in order to compute the output probabilities.

Since the RNN was bidirectional, no output delay was added during training. To train the RNN, data from all available sessions were used. Instead of training and testing on separate blocks of data, as was done for the real-time performance evaluation, we used all blocks of data (both the "open-loop" blocks where no real-time decoding occurred, and the "closed-loop" blocks with real-time decoding). We randomly selected 10% of the sentences from each day as held-out test sentences for evaluation. We excluded all blocks of the 7 repeated sentences that we collected for comparison with (Pandarinath et al., 2017), since we didn't want the RNN to overfit to these sentences.

We used the following training parameters: $\lambda$=1e-5, H=512, $\sigma$=1.2, $\gamma$=0, $\zeta$=0, $\chi$=0.375, $\omega$=0.1, $\alpha$=0.01, M=100k, N=64, T=24.

#### RNN Parameter Sweeps and Variants

Retrospectively, we tested the effect of five important parameters on RNN performance (SFig. 3): (1) the amount of synthetic data used during training, (2) the amount of artificial white noise added to the inputs during training, (3) the amount of artificial feature mean noise added during training, (4) whether the RNN was trained with all days of data or just a single day, and (5) whether multiple days were combined with separate input layers or the same input layer. The results confirm that synthetic data, input white noise, feature mean noise, and combining data across days with separate input layers were all essential for high performance

When training multi-day RNNs for this analysis, we used all 10 sessions where open-loop data was collected (i.e., blocks where T5 was handwriting sentences but no real-time decoding was performed). We then evaluated performance on the 7 sessions that had both open-loop data and closed-loop copy typing data. We evaluated the character error rate on either held-out open-loop sentences ("held-out trials") or held-out closed-loop blocks ("held-out blocks") which occurred later in the session (~1 hour later). For the held-out trials, we held out 10% of the open-loop trials at random; for the held-out blocks, all data were used. When training single-day RNNs that used only a single session of data, we trained separate RNNs for all 7 evaluation sessions.

For testing the effect of synthetic data and input white noise (SFig. 3A), we performed a joint grid search over both the synthetic data fraction and amount of white noise. Since both parameters have a regularizing effect, we searched over both at the same time so we could make more definitive statements about whether both were really needed (or whether just one tuned to the correct value provided enough regularization to reach peak performance). We searched over four possible values of the synthetic data fraction (0, 0.25, 0.5, and 0.75) and five possible values of white noise standard deviation (0, 0.6, 1.2, 1.8, and 2.4), making for a 4 x 5 grid. The character error rates shown in SFig. 3A were averaged over the 7 evaluation days.

For testing the effect of feature mean noise (SFig. 3B), results are shown for each of four pairs of constant offset noise ($\gamma$) and random walk noise ($\zeta$): ($\gamma$=0, $\zeta$=0), ($\gamma$=0.3, $\zeta$=0.01), ($\gamma$=0.6, $\zeta$=0.02), ($\gamma$=1.2, $\zeta$=0.04). For each pair, 3 separate RNNs were trained.

For testing the effect of training on all days of data vs. just a single day (SFig. 3C), results are shown from one multi-day RNN and seven single-day RNNs (one for each evaluation session).

For testing the effect of using separate input layers for each day vs. a single layer when training across multiple days (SFig. 3D), results are shown from one separate-layer RNN and one shared-layer RNN.

### Comparison to a Hidden Markov Model Decoder

To test whether an RNN was necessary for achieving high-performance compared to a simpler decoding approach, we tested the performance of a straightforward hidden Markov model decoder (Table S4). The results confirm that our RNN strongly outperforms a simple HMM, especially for held-out blocks where the feature means are likely to have changed substantially due to neural non-stationarities (Jarosiewicz et al., 2015; Downey et al., 2018).

We designed the HMM decoder in the same way as we designed the forced-alignment HMMs used to label the sentence data, except instead of containing character states that marched forward through a fixed sequence of characters, each character could transition to any other character with equal probability. We also tweaked the state transition probabilities within each character to the following values (to improve decoding performance):

$$P(s_{x,j} \rightarrow s_{x,j}) = 0.4$$
$$P(s_{x,j} \rightarrow s_{x+1,j}) = 0.2$$
$$P(s_{x,j} \rightarrow s_{x+2,j}) = 0.4$$

When evaluating decoder performance, we applied the language model to both the RNN and the HMM. Using a language model makes the comparison fairer by compensating for the fact that the RNN itself could learn character transition probabilities that better model the English language than the simple uniform transition model we used for the HMM.

#### Alphabet Optimization

We solved the following optimization problem to find a new set of 26 letters that maximizes the distance between the closest pair of letters (SFig. 4A):

$$\underset{X_1, X_2, \ldots, X_{26}}{\operatorname{argmax}} \min_{i,j} \left\| S(Q(X_i)) - S(Q(X_j)) \right\|_F^2$$

Here, $X_i$ is a 2 x 100 matrix that represents the pen tip velocity trajectory for letter *i* (each column is a velocity vector for one time step), *Q* is a “squashing” function that constrains each column of $X_i$ to lie within the unit circle, *S* is a smoothing function that constrains the trajectories to be smooth by convolving each row with a Gaussian kernel (sd = 8), and $\|X\|_F$ is the Frobenius norm of X (i.e., the square root of the sum of squared entries in the matrix X).

*Q* was defined as follows (where $x_i$ is the *i*th column of *X*):

$$Q(X) = [q(x_1) \quad q(x_2) \quad \ldots \quad q(x_{100})]$$
$$q(x_i) = \frac{x_i}{\|x_i\|}\left(1 - e^{-\|x_i\|}\right)$$

We found local minima of the above cost function by using gradient descent (implemented with TensorFlow v1.15 to compute the gradients, using the “Adam” optimization method (Abadi et al., 2016; Kingma and Ba, 2017)).

We varied the width of the Gaussian kernel used for smoothing until the temporal dimensionality of the optimized letters was equal to the dimensionality of the Latin alphabet (~4D), resulting in a set of letters that has a visually similar level of complexity and curviness to the Latin letters (SFig. 4A). To define the pen trajectories of the Latin alphabet, we used the trajectories reconstructed in Figure 1, which capture how T5 wrote the letters. These trajectories were resampled (linearly time-warped) to 100 time steps (as was done in Figure 3).

As a comparison, we also optimized for a set of 26 straight lines (SFig. 4B). To do so, we used the above cost function except that each column of $X_i$ was constrained to be equal, enforcing a straight-line trajectory.

Note that no neural encoding model is explicitly considered in the optimization. However, if we assume linear neural tuning to pen tip velocity (i.e., “cosine tuning”), then the cost functions are equivalent under reasonable assumptions of uniformity in the neural tuning. That is, maximizing the nearest neighbor distance of pen trajectories is the same as maximizing the nearest neighbor distance of evoked neural features under this assumption. The result follows from the fact that distances are preserved under orthogonal transformations.

To show this, let *E* be a matrix of linear tuning coefficients (of size 192 x 2) for 192 hypothetical neural features. Assume the neural features are tuned to pen tip velocity in the following way:

$$f_t = Ev_t + b$$

Here, $f_t$ is a 192 x 1 vector of neural features for time step $t$, $v_t$ is a 2 x 1 pen tip velocity vector, and $b$ is a 192 x 1 offset vector. The squared distance between the neural feature matrices associated with two different letters can then be expressed in the following way (where $v_t$ is the pen tip velocity for letter A and $u_t$ is the pen tip velocity for letter B):

$$\sum_{t=1}^{N} (Ev_t + b - [Eu_t + b])^T \, (Ev_t + b - [Eu_t + b])$$

$$= \sum_{t=1}^{N} (E(v_t - u_t))^T \, (E(v_t - u_t))$$

$$= \sum_{t=1}^{N} (v_t - u_t)^T E^T E \, (v_t - u_t)$$

Let us assume that the columns of $E$ are roughly orthogonal to each other, which would be the case for "uniform" neural tuning where the rows of $E$ (i.e., the "preferred directions") are uniformly distributed about the origin. Then, $E^TE$ is a diagonal matrix with approximately equal entries along the diagonal (we can denote this entry as $\alpha$). Then, we have:

$$\sum_{t=1}^{N} (v_t - u_t)^T E^T E \, (v_t - u_t)$$

$$\approx \sum_{t=1}^{N} (v_t - u_t)^T \begin{bmatrix} \alpha & 0 \\ 0 & \alpha \end{bmatrix} (v_t - u_t)$$

$$= \alpha \sum_{t=1}^{N} (v_t - u_t)^T \, (v_t - u_t)$$

Thus, linear neural tuning to pen tip velocity implies that the neural distances are directly proportional to the distances between the velocity trajectories themselves.

More generally, the neural encoding of a pen trajectory is not solely linear. However, we conjecture that as long as the neural encoding function is a reasonably smooth function of the pen tip velocity, then far-apart pen trajectories are likely to evoke far-apart neural activity, making it a reasonable approach to optimize over pen trajectories directly.

### Pen Trajectory Visualization

We trained the pen velocity decoder in an iterative process, where the templates were also shifted and time-warped to maximize decoding performance (since drawing the characters with a mouse may not yield the same writing speed and reaction times as T5). Three iterations were performed. Each iteration began with training the linear decoder with the current templates (using ordinary least squares regression to minimize the error between the template velocities and the decoded velocities). Then, the template start times and time dilations were optimized (via a grid search) to best match the decoded velocities (maximize the correlation). Finally, to visualize the character shapes, the decoder was applied to the time-warped and trial-averaged neural activity. The decoded velocity was integrated to yield the pen tip position.

Specifically, each trial of neural activity was represented by a matrix of binned and smoothed threshold crossing rates with dimensions 200 x 192 (200 time steps and 192 electrodes) taken from a -0.5 to 1.5 second window around the go cue. Then, these matrices were concatenated vertically to form a predictor matrix for the linear regression of size 200*N x 192, where N is the number of trials. Finally, a column of ones was added to the predictor matrix (to create a constant offset term), yielding a design matrix $X$ of dimension 200*N x 193. We also constructed a response matrix $Y$ of dimension 200*N x 2 containing the template velocity vectors at each time step that the decoder should predict.

For the first iteration, $Y$ was constructed by setting the entries of each trial equal to the velocity vectors in that character's template. The decoder was then computed with ordinary least squares to minimize the following mean squared error cost function:

$$\|Y - XB\|_F^2$$

For the subsequent two iterations, we used the decoded velocity vectors $XB$ to time-warp and shift the character templates before constructing $Y$. Specifically, we performed a grid search for each character, searching for possible template shift times between -0.4 and 0.4 seconds and possible template warping factors between 0.5 and 2.0. Templates were time-warped by resampling with linear interpolation. The shift time and warp factor that maximized the correlation between the template and the previously decoded velocity vectors was chosen. Finally, the shifted and warped templates were used to create a new $Y$ matrix, with which a new decoder could be built.

### References

Abadi, M., Agarwal, A., Barham, P., Brevdo, E., Chen, Z., Citro, C., Corrado, G.S., Davis, A., Dean, J., Devin, M., et al. (2016). TensorFlow: Large-Scale Machine Learning on Heterogeneous Distributed Systems. ArXiv:1603.04467 [Cs].

Brants, T., and Franz, A. (2006). Web 1T 5-gram Version 1.

Degenhart, A.D., Bishop, W.E., Oby, E.R., Tyler-Kabara, E.C., Chase, S.M., Batista, A.P., and Yu, B.M. (2020). Stabilization of a brain–computer interface via the alignment of low-dimensional spaces of neural activity. Nature Biomedical Engineering 1–14.

Downey, J.E., Schwed, N., Chase, S.M., Schwartz, A.B., and Collinger, J.L. (2018). Intracortical recording stability in human brain–computer interface users. J. Neural Eng. *15*, 046016.

Goodfellow, I., Bengio, Y., and Courville, A. (2016). Deep Learning (The MIT Press).

Graves, A., Fernández, S., Gomez, F., and Schmidhuber, J. (2006). Connectionist temporal classification: labelling unsegmented sequence data with recurrent neural networks. In Proceedings of the 23rd International Conference on Machine Learning, (Pittsburgh, Pennsylvania, USA: Association for Computing Machinery), pp. 369–376.

Jarosiewicz, B., Sarma, A.A., Bacher, D., Masse, N.Y., Simeral, J.D., Sorice, B., Oakley, E.M., Blabe, C., Pandarinath, C., Gilja, V., et al. (2015). Virtual typing by people with tetraplegia using a self-calibrating intracortical brain-computer interface. Science Translational Medicine *7*, 313ra179-313ra179.

Kingma, D.P., and Ba, J. (2017). Adam: A Method for Stochastic Optimization. ArXiv:1412.6980 [Cs].

Mohri, M., Pereira, F., and Riley, M. (2008). Speech Recognition with Weighted Finite-State Transducers. In Springer Handbook of Speech Processing, J. Benesty, M.M. Sondhi, and Y.A. Huang, eds. (Berlin, Heidelberg: Springer), pp. 559–584.

Pandarinath, C., Nuyujukian, P., Blabe, C.H., Sorice, B.L., Saab, J., Willett, F.R., Hochberg, L.R., Shenoy, K.V., and Henderson, J.M. (2017). High performance communication by people with paralysis using an intracortical brain-computer interface. ELife *6*, e18554.

Poole, B., Williams, A.H., Maheswaranathan, N., Yu, B., Santhanam, G., Ryu, S.I., Baccus, S., Shenoy, K.V., and Ganguli, S. (2017). Time-warped PCA: simultaneous alignment and dimensionality reduction of neural data. Cosyne Abstracts.

Povey, D., Ghoshal, A., Boulianne, G., Burget, L., Glembek, O., Goel, N., Hannemann, M., Motlicek, P., Qian, Y., Schwarz, P., et al. (2011). The Kaldi Speech Recognition Toolkit. IEEE 2011 Workshop on Automatic Speech Recognition and Understanding.

Puigcerver, J. (2017). Are Multidimensional Recurrent Layers Really Necessary for Handwritten Text Recognition? In 2017 14th IAPR International Conference on Document Analysis and Recognition (ICDAR), pp. 67–72.

Rabiner, L.R. (1989). A tutorial on hidden Markov models and selected applications in speech recognition. Proceedings of the IEEE *77*, 257–286.

Radford, A., Wu, J., Child, R., Luan, D., Amodei, D., and Sutskever, I. (2018). Language models are unsupervised multitask learners. OpenAI Technical Report.

Stavisky, S.D., Willett, F.R., Wilson, G.H., Murphy, B.A., Rezaii, P., Avansino, D.T., Memberg, W.D., Miller, J.P., Kirsch, R.F., Hochberg, L.R., et al. (2019). Neural ensemble dynamics in dorsal motor cortex during speech in people with paralysis. ELife *8*, e46015.

Stolcke, A., Zheng, J., Wang, W., and Abrash, V. (2011). SRILM at Sixteen: Update and Outlook.

Sussillo, D., Stavisky, S.D., Kao, J.C., Ryu, S.I., and Shenoy, K.V. (2016). Making brain–machine interfaces robust to future neural variability. Nat Commun *7*.

Willett, F.R., Deo, D.R., Avansino, D.T., Rezaii, P., Hochberg, L.R., Henderson, J.M., and Shenoy, K.V. (2020). Hand Knob Area of Premotor Cortex Represents the Whole Body in a Compositional Way. Cell.

Williams, A.H., Poole, B., Maheswaranathan, N., Dhawale, A.K., Fisher, T., Wilson, C.D., Brann, D.H., Trautmann, E.M., Ryu, S., Shusterman, R., et al. (2020). Discovering Precise Temporal Patterns in Large-Scale Neural Recordings through Robust and Interpretable Time Warping. Neuron *105*, 246-259.e8.

Xiong, W., Wu, L., Alleva, F., Droppo, J., Huang, X., and Stolcke, A. (2017). The Microsoft 2017 Conversational Speech Recognition System. ArXiv:1708.06073 [Cs].

Young, S.J., Evermann, G., Gales, M.J.F., Kershaw, D., Moore, G., Odell, J.J., Ollason, D.G., Povey, D., Valtchev, V., and Woodland, P.C. (2006). The HTK book version 3.4.
